# *Cis*-regulatory control of transcriptional timing and noise in response to estrogen

**DOI:** 10.1101/2023.03.14.532457

**Authors:** Matthew Ginley-Hidinger, Hosiana Abewe, Kyle Osborne, Alexandra Richey, Noel Kitchen, Katelyn L. Mortenson, Erin M. Wissink, John Lis, Xiaoyang Zhang, Jason Gertz

**Affiliations:** Huntsman Cancer Institute, University of Utah, Salt Lake City, UT 84112, USA; Department of Biomedical Engineering, University of Utah, Salt Lake City, UT 84112, USA; Department of Oncological Sciences, University of Utah, Salt Lake City, UT 84112, USA; Department of Molecular Biology and Genetics, Cornell University, Ithaca, NY 14853, USA

## Abstract

Cis-regulatory elements control transcription levels, temporal dynamics, and cell-cell variation or transcriptional noise. However, the combination of regulatory features that control these different attributes is not fully understood. Here, we used single cell RNA-seq during an estrogen treatment time course and machine learning to identify predictors of expression timing and noise. We find that genes with multiple active enhancers exhibit faster temporal responses. We verified this finding by showing that manipulation of enhancer activity changes the temporal response of estrogen target genes. Analysis of transcriptional noise uncovered a relationship between promoter and enhancer activity, with active promoters associated with low noise and active enhancers linked to high noise. Finally, we observed that co-expression across single cells is an emergent property associated with chromatin looping, timing, and noise. Overall, our results indicate a fundamental tradeoff between a gene’s ability to quickly respond to incoming signals and maintain low variation across cells.

## Introduction

Cis-Regulatory elements (CREs) control the precise spatiotemporal expression of genes across the genome. In addition to a gene’s promoter, many enhancers collaborate to control a single gene’s expression in mammalian cells (ENCODE, 2012; Kundaje et al., 2015; Zhang et al., 2020). External chemical signals often induce changes in cell phenotypes by altering transcription, requiring coordinated gene expression programs. Signal transduction can lead to transcription factor (TF) binding changes and epigenetic modifications at CREs (MacKenzie et al., 2013). For cells to appropriately respond to stimuli, CREs must guide the amount of transcript produced (MacKenzie et al., 2013), the timing of transcriptional changes (Kolch et al., 2015; Wei et al., 2016), and the amount of transcriptional variation or noise (Kolch et al., 2015; Raj and Van Oudenaarden, 2008; Raser and O’Shea, 2005). While there has been extensive research on the role that CREs play in transcription levels, less is understood about the properties of CREs that control gene expression timing and noise.

Temporal regulation of gene expression is an essential attribute of transcriptional control for cellular processes such as cell fate transitions (Basma et al., 2009; Chamberlain et al., 2008; Konstantinides et al., 2022) and responses to signals (Behar and Hoffmann, 2010; Krakauer et al., 2002; Uribe et al., 2021). Specific genes, often termed immediate-early genes, are rapidly activated in response to a signal, while other genes change expression more gradually (Sheng and Greenberg, 1990; Uhlitz et al., 2017; Uribe et al., 2021). Genes that show coordinated trajectories are often functionally related, driving diverse phenotypes at different timescales (Gandhi et al., 2011; Krakauer et al., 2002; Schnoes et al., 2008; Szustakowski et al., 2007). Previous studies have identified several mechanisms that regulate transcriptional timing. One influential factor is the state of a gene’s promoter. For example, pre-loading of RNA polymerase II (RNAPII) at the promoter is indicative of earlier gene expression responses (Tullai et al., 2007). Additional promoter features associated with early responding genes include TATA motifs at the promoter, a greater number of TF binding motifs, and increased chromatin accessibility (Murai et al., 2020; Tullai et al., 2007). Enhancers are also crucial for gene expression timing. Inhibition or deletion of specific enhancers can prolong the time needed for a gene to reach maximal expression without altering final expression levels (Juan and Ruddle, 2003; Simeonov et al., 2017). Stretches of potent enhancers, called super-enhancers, regulate some immediate-early genes (Hah et al., 2015). In contrast, enhancers marked by repressive chromatin marks, termed latent enhancers, exhibit slower activation and are associated with late-responding genes (Ostuni et al., 2013). Overall, relatively little is known about which genomic features in a gene’s cis-regulatory repertoire are important for influencing stimulus-dependent temporal gene expression responses.

In addition to regulating gene expression timing and levels, CREs control the amount of transcriptional noise. Transcriptional noise is a combination of intrinsic stochasticity and extrinsic variability that cause transcript variation across a population of isogenic cells (Elowitz et al., 2002; Fraser et al., 2021b; Kundaje et al., 2015). Cells must regulate transcriptional variation, as both high and low variation have functional consequences. High variation can have benefits, as cells may be more adaptable to changing environments (Pedraza et al., 2018; Wollman, 2018) and more likely to undergo cell fate transitions (Desai et al., 2021; Suderman et al., 2017). Noise may additionally confer the ability of a cell population to produce a diverse output to a single incoming signal (Azpeitia et al., 2020). However, noise can be associated with negative consequences, such as worse cancer outcomes (Han et al., 2016), cancer therapy resistance (Qin et al., 2020; Shaffer et al., 2017), and the ability of cancer cells to metastasize (Fidler, 1978; Nguyen et al., 2016). Both promoters and enhancers can regulate intrinsic noise kinetics and sensitivity to extrinsic noise sources (Larsson et al., 2019). For example, nucleosome positioning and histone modifications at the promoter are important noise regulators (Choi and Kim, 2009; Dadiani et al., 2013; Fraser et al., 2021b; Nicolas et al., 2018; Wu et al., 2017), with active histone marks at promoters often associated with low noise (Urban and Johnston, 2018). Additionally, a greater number of transcription factors binding at a promoter may be a basis for greater amounts of noise (Parab et al., 2022). The role of enhancers in controlling mammalian expression noise is less clear. Thermodynamic modeling approaches suggest that multiple enhancers should buffer noise (Hnisz et al., 2017), while experimental evidence shows that super-enhancers are generally associated with noisier expression (Fraser et al., 2021b; Wibisana et al., 2022). A remaining challenge is understanding the effects of multiple enhancers in combination with a promoter on expression noise.

To investigate the regulatory control of timing and noise in depth, we focused on the transcriptional response to estrogens. Estrogen Receptor Ill (ER) is a nuclear hormone receptor activated by estrogens, including endogenously produced 17β-estradiol (E2). In the presence of E2, ER becomes an active TF and regulates the expression of hundreds of genes (Bjornstrom and Sjoberg, 2005). ER is a clinically relevant TF, a potent oncogenic driver for endometrial and breast cancer (Rodriguez et al., 2019a; Stanford et al., 1986), and a well-studied model TF. Upon activation, ER both upregulates and downregulates genes at different timescales (Frasor et al., 2003; Liberzon et al., 2015). Following an estrogen induction, ER activates successive sets of functionally unique genes, as seen in genes related to vascularization, signaling, proliferation, and cell cycle (Jagannathan and Robinson-Rechavi, 2011; Schnoes et al., 2008). ER has also been shown to regulate transcriptional noise. Live cell imaging of ER target genes *GREB1* (Fritzsch et al., 2018) and *TFF1* (Rodriguez et al., 2019b) show that ER impacts transcriptional noise by modulating transcription kinetics. The temporal, heterogeneous complexity of the ER transcriptional program makes it an ideal model system for studying how CREs regulate transcriptional timing and noise in response to an external stimulus.

To better understand the genomic underpinnings of transcriptional levels, timing, and noise, we analyzed the transcriptional response to E2 using a time course of single cell RNA-seq (scRNA-seq) in two cell types (human breast and endometrial cancer cells). Feature ranking approaches, using genomic data, revealed important determinants that control these transcriptional attributes. A strong enhancer repertoire was associated with earlier changes in gene expression, which was confirmed using functional perturbation by dCas9-based synthetic transcription factors. Promoter features also regulate timing, such as transcriptional repressor SIN3A being found at the promoters of “Late” genes. We uncovered a balance between enhancers and promoters in regulating expression noise, where strong enhancers drive higher noise and strong promoters are associated with low expression variance. The role of enhancers in timing and noise reveals a tradeoff between expression noise and the ability to respond quickly to incoming signals.

## Results

### Machine learning approach accurately predicts genomic determinants of expression levels

To uncover features of gene regulation that control expression levels, timing, and noise, pooled scRNA-seq was conducted following 0-, 2-, 4-, and 8-hour E2 treatments in two cell lines: Ishikawa (human endometrial adenocarcinoma) and T-47D (human breast carcinoma) (QC metrics prior to filtering cells shown in Figure S1A and B). After filtering very low expressed genes based on mean expression, we observed 12,756 and 11,395 genes across all time points in Ishikawa and T-47D cells, respectively. We first set out to identify determinants of mean expression levels by focusing on the 0-hour timepoint (no E2 treatment). Genome-wide data, mostly on protein-DNA interactions, from publicly available sources (ENCODE, 2012; Shu et al., 2016; Zhang et al., 2016) and experiments conducted for this study (Table S3) were quantified at promoter and enhancer regions. Due to variations in enhancer number and strength across genes, an aggregate enhancer score was used to capture the combined action of multiple enhancers (see Methods) (Figure S1C). Genomic features were ranked by importance for classifying low (bottom 20% of genes), medium (middle 60%), and high (top 20% of genes) expression levels using the Boruta algorithm for feature selection (Kursa and Rudnicki, 2010a), which has been previously used to uncover determinants of expression noise in *drosophila* (Sigalova et al., 2020). For feature ranking, we grouped genes from both cell types to find mutual predictors, with the expectation that there are common underlying mechanisms for transcription control.

Elements of the pre-initiation complex and H3K27ac at the promoter ranked as the most important predictors for transcript levels (Figure 1A left). These features had stronger promoter signals at higher expressed genes (Figure 1B-E), in agreement with previous literature (Schier and Taatjes, 2020; Wang et al., 2008). Important factors at promoters and enhancers showed a general trend of increased signal for highly expressed genes (Figure 1A, right). Our dataset is strongly biased toward activating transcription factors and histone modifications found at active regulatory regions. Plotting the average promoter intensity compared to the average enhancer score across all confirmed datasets verifies that strong promoters and active enhancers are associated with higher gene expression levels (Figure 1F). Overall, the Boruta approach was successful at identifying known predictors of transcript levels.

**Figure 1.**
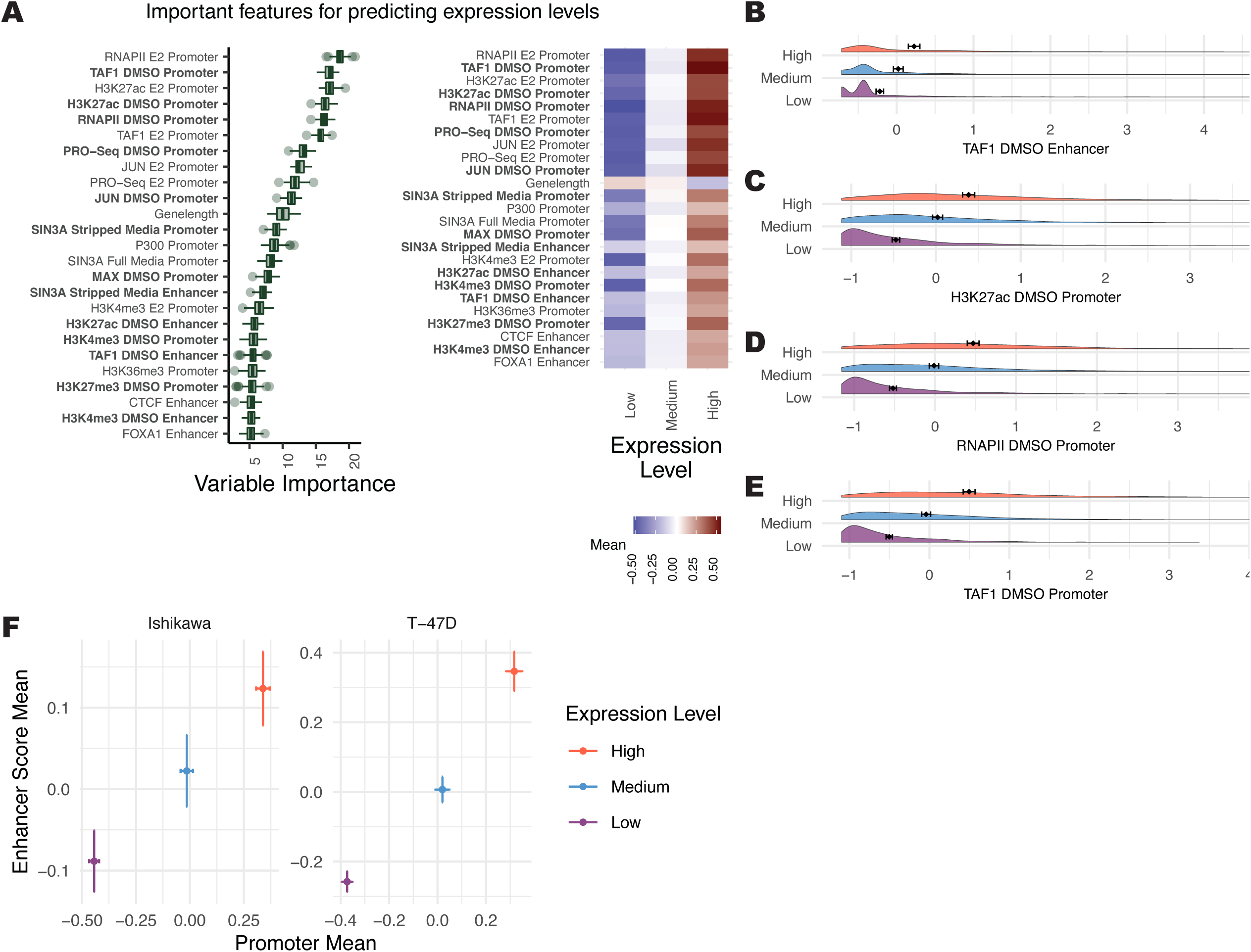
Several genomic features associate with gene expression levels. (A, left) Boruta feature ranking of genomic features shows importance of a feature for predicting mean levels. (A, right) Average signal intensity for each genomic dataset, grouped by mean expression levels, is shown. Datasets shown in bold were performed in the absence of ER activation. (B-E) Distributions of the top 4 most important ranked features in the absence of ER activation, separated by mean expression levels, show higher signal for “High” expression groups. X-axis represents Z-scores and error bars show the mean ± 95% confidence intervals. (F) Mean enhancer score signal for all Boruta confirmed features vs. mean promoter signal across all confirmed features is shown where error bars show the mean ± 95% confidence intervals. Axes do not represent the full range.

### Analysis of temporal trajectories indicates features that control estrogen response timing

We next analyzed genomic predictors of response timing using the scRNA-seq data following 0-, 2-, 4-and 8-hours of estrogen treatment. Using dimensionality-reduction UMAP plots, the temporal progression of the E2 response in single cells can be observed, although cells are not fully separated by timepoint (Figure S1D and E). The clustering of cells suggest that the transcriptional response does not follow a tight temporal pattern and that variation in the E2 response exists between cells. In addition, T-47D cells exhibit distinct clusters prior to E2 treatment that do not appear to be cell cycle related based on *TOP2A* expression (Figure S1F). Based on a Wilcoxon rank-sum test (Bauer, 1972), there were between 491 and 1146 differentially expressed genes for each cell type between any E2 treatment timepoint and the 0-hour control (Figure S2A). scRNA-seq summed counts showed high concordance with previously published bulk RNA-seq data in the same cell lines (Figure S2B and C) (Gertz et al., 2013). Compared to bulk RNA-seq, there are more differentially expressed genes with high expression (Figure S2B and C) and lower fold changes (Figure S2D and E), likely due to the increased statistical power of scRNA-seq for calling differential expression of highly expressed genes. However, there was still high overlap between differentially expressed genes at the matching 0-hour vs. 8-hour comparison. Of genes that occur in both single cell and bulk datasets 48.1% and 73.6% of 8-hour bulk RNA-seq genes overlap with 8-hour scRNA-seq genes (p <2.2×10^-16^, odds ratio = 6.11; and p < 2.2×10^-16^, odds ratio=12.61; fisher’s exact test in Ishikawa and T-47D, respectively). In addition to the comparison with previously published bulk RNA-seq from 0-and 8-hour E2 treatment, we performed bulk RNA-seq using the same time course as the scRNA-seq experiment. We observed highly significant overlaps between expression trajectories in both cell lines (Figure S2F). scRNA-seq can therefore be a valuable tool to capture subtle changes in gene expression following E2 treatment.

Based on the timepoint at which a gene is differentially regulated, genes were classified into temporal response trajectories for up-and down-regulated genes. One class of genes rapidly changes expression in response to E2 (termed Early genes), while another class changes more gradually and takes longer to reach a maximum response (termed Late genes). A representative set of Control genes was also randomly selected using stratified sampling to mirror mean levels found in differential genes. Early responding genes have a significant initial response to E2 by 2 hours, then return toward baseline for both up-and down-regulation, consistent with previous reports of pulse-like expression in immediate-early genes (Iyer et al., 1999) (Figure 2A and B). In contrast, genes classified as Late show a slow and steady response over time (Figure 2A and B). We found that both Early Up and Late Up genes were significantly enriched for the previously described hallmarks of estrogen response early and late (Figure S3A-D). As we have reported previously (Gertz et al., 2012; Gertz et al., 2013), the transcriptional response to E2 is highly cell type-specific. However, genes with Up trajectories are more conserved between cell types than genes with Down trajectories (Figure S3E) (p-value=0.013; t-test).

**Figure 2.**
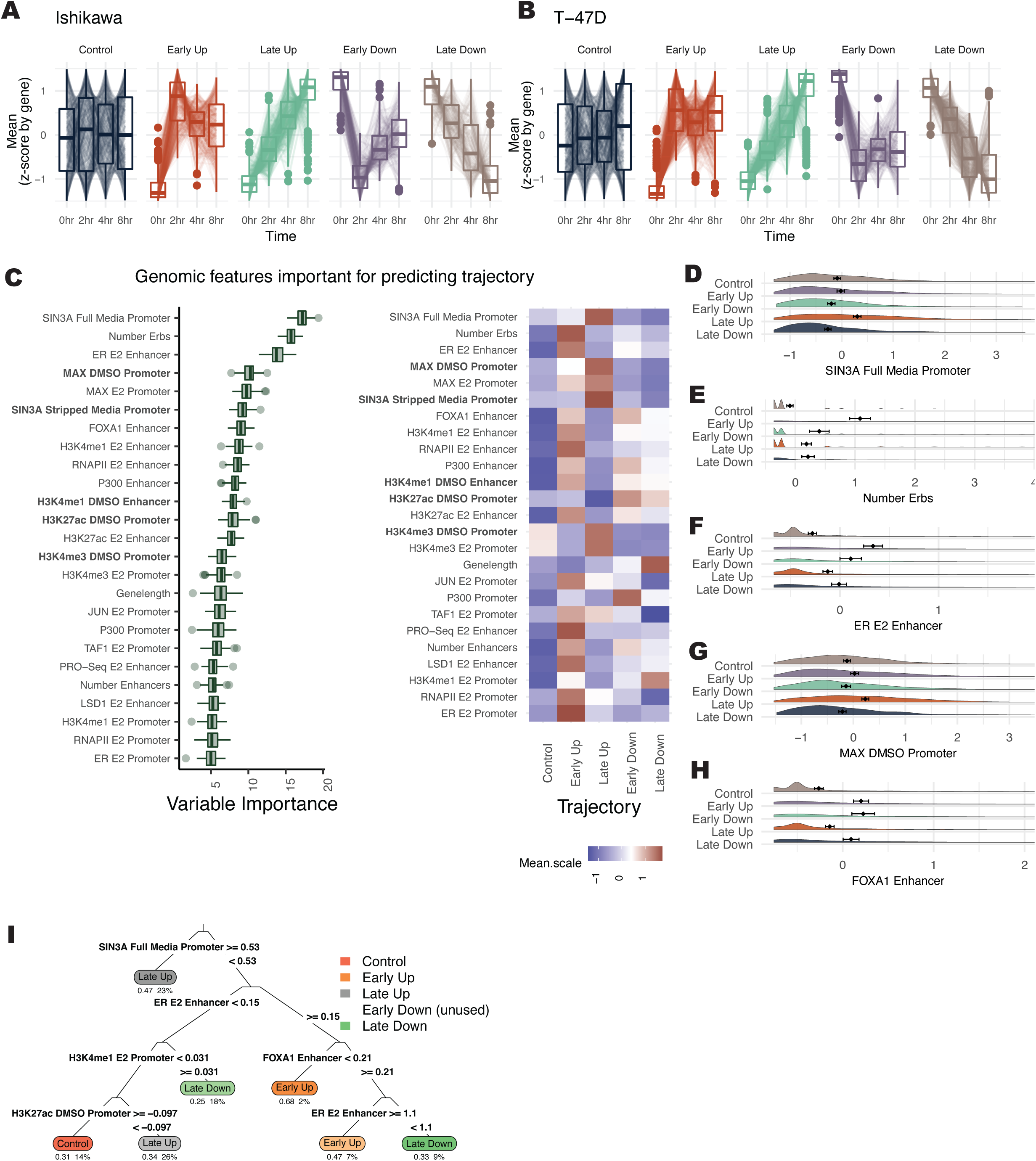
SIN3A and multiple ER bound sites are the strongest predictors of transcriptional response timing. (A-B) Z-scores for each gene across 4 timepoints are shown within different gene expression trajectories in (A) Ishikawa and (B) T-47D cells. (C, left) Based on Boruta ranking, the top 25 most important features are shown for classifying gene trajectories. (C, right) Heatmap displays the average signal by trajectory for each predictor. Datasets shown in bold were performed in the absence of ER activation. (D-H) Distribution of signal (Z-score) of the most important features for predicting temporal trajectories is shown. (I) Decision tree shows the hierarchy of classification for predicting gene expression trajectories. The top 4 layers are shown.

The Boruta algorithm was used to identify predictors of temporal trajectories. SIN3A signal at the promoter was most predictive of gene expression trajectory and is associated with Late Up genes (Figure 2C and D). MAX, which is known to repress genes through recruitment of SIN3A (Baudino and Cleveland, 2001), was also classified as important and enriched at promoters of Late Up genes (Figure 2G). The number of ER binding sites (ERBS) that loop to the promoter and the ER signal at enhancers were the next most important features (Figure 2C). These two features are enriched in Early Up genes (Figure 2E and F). A higher number of total enhancers is enriched for genes that respond Early (Figure S3F); however, specific proteins (e.g., FOXA1) (Figure 2H) are more balanced between Early Up and Early Down genes than other factors (e.g., ER) that show preferential binding near Early Up genes. Together, these results suggest that the number of enhancers plays a prominent role in the temporal response of genes, but transcription factors at these sites, such as ER, help control the direction and timing of gene expression changes. An example of an optimal decision tree was computed to examine a potential hierarchy of factors determining a gene’s temporal response (Figure 2I). This decision tree shows how SIN3A is the primary separator of genes into the Late Up trajectory, which may take precedence over ER signal. This analysis was run on the combined Ishikawa and T-47D data. When we perform the same analysis separately on each cell line, we find that the importance scores are highly correlated (r = 0.69; Figure S3G), indicating that shared mechanisms control timing across cell types. When analyzing promoter and enhancer activity separately, we found that promoter signal is highly correlated between cell types regardless of whether or not a gene exhibited the same response to E2 (r = 0.74 vs. 0.7). However, enhancer signal is less correlated between cell types and lower when a gene’s trajectory is not conserved between cell types (r = 0.39 vs. 0.3), indicating that enhancer features play a larger role in cell type-specific gene regulation by estrogen. Overall, Boruta analysis of temporal trajectories uncovers unique factors that may regulate transcription response timing and shows the association of multiple ER bound enhancers with a rapid up-regulation in response to E2.

One potential confounding factor of the trajectory analysis is the impact of mRNA half-life, since the scRNA-seq measurements represent poly-adenylated mRNAs and not simply nascent transcripts. To evaluate how mRNA turnover could play a role in trajectory, we compared previously published mRNA half-life measurements (Agarwal and Kelley, 2022) across E2 regulated genes (Figure S3H). We found that Early Down genes exhibited significantly shorter mRNA half-lives than other genes (p < 2.2×10^-16^, Kolmogorov-Smirnov test; average of 0.55 standard deviations below control genes). This finding suggests that fast mRNA turnover is important for quick down-regulation in response to estrogen treatment. To a lesser extent, we see that Early Up genes are also characterized by shorter half-lives (p = 8.×10^-12^, Kolmogorov-Smirnov test; average of 0.29 standard deviations below control genes). This feature may help to explain the pattern of Early Up genes rapidly increasing by 2 hours, but leveling off during the rest of the 8-hour E2 treatment.

### Functional perturbation of CREs alters temporal responses

To test the functional relationship between CREs and transcriptional response timing, dCas9-based activators and repressors were used to modulate the genomic activity of regulatory regions in Ishikawa cells, as T-47D cells exhibit low transfection efficiencies. Gene expression responses were then measured during a time course of E2 treatment using quantitative PCR. A SID(4x)-dCas9-KRAB construct was used for repression (Carleton et al., 2017). This construct can directly recruit SIN3A, a good predictor of Late gene expression responses. It also recruits Histone deacetylases (HDACs) (Urrutia, 2003), corresponding to the low H3K27ac seen at Late Up genes. For activation, dCas9-VP16(10x) was used. We have previously shown that dCas9-VP16(10x) modulates expression from enhancers and induces acetylation at targeted regions (Ginley-Hidinger et al., 2019). dCas9-VP16(10x) recruits many activating cofactors, including members of the pre-initiation complex and p300, which are associated with Early genes.

*TACSTD2* is an E2 regulated gene that is a prognostic indicator for endometrial cancer disease-free survival (Bignotti et al., 2012), is overexpressed in some breast cancers (Shvartsur and Bonavida, 2014), and exhibits an Early Up trajectory. Targeting the enhancers of *TACSTD2*, marked by H3K27ac and ER binding (Figure 3A) with SID(4X)-dCas9-KRAB resulted in a slower, more gradual response to E2. The time for expression to reach maximal observed levels was increased when targeting 2 out of 3 individual enhancers (Figure 3B). When targeting enhancer +4.7kb and enhancer −15.2kb, longer activation times were observed compared to non-targeting controls based on time to half maximal expression (Figure 3D). Enhancer +4.7kb and enhancer −15.2kb exhibited slower activation rates, but reached similar activation levels at 8 hours compared to controls, indicating that these enhancers can regulate the timing of a response without affecting overall levels. Additionally, the slope of gene expression over time showed generally slower initial activation when *TACSTD2* regulatory regions were inhibited compared to the control, followed by more similar rates at later timepoints (Figure 3F). Synthetic activation of the same *TACSTD2*-linked CREs led to a more rapid response when targeting most individual enhancers and the combination of all enhancers (Figure 3C), as evidenced by decreased time to half maximal expression (Figure 3E). Again, we see that enhancer −15.2kb changes expression timing without affecting overall transcript abundance. Analysis of the slopes revealed a generally increased initial slope with respect to the control when *TACSTD2* regulatory regions were targeted (Figure 3G). These results imply that decreasing enhancer activity can reduce initial activation rates, while activating enhancers can potentiate quicker responses to E2.

**Figure 3.**
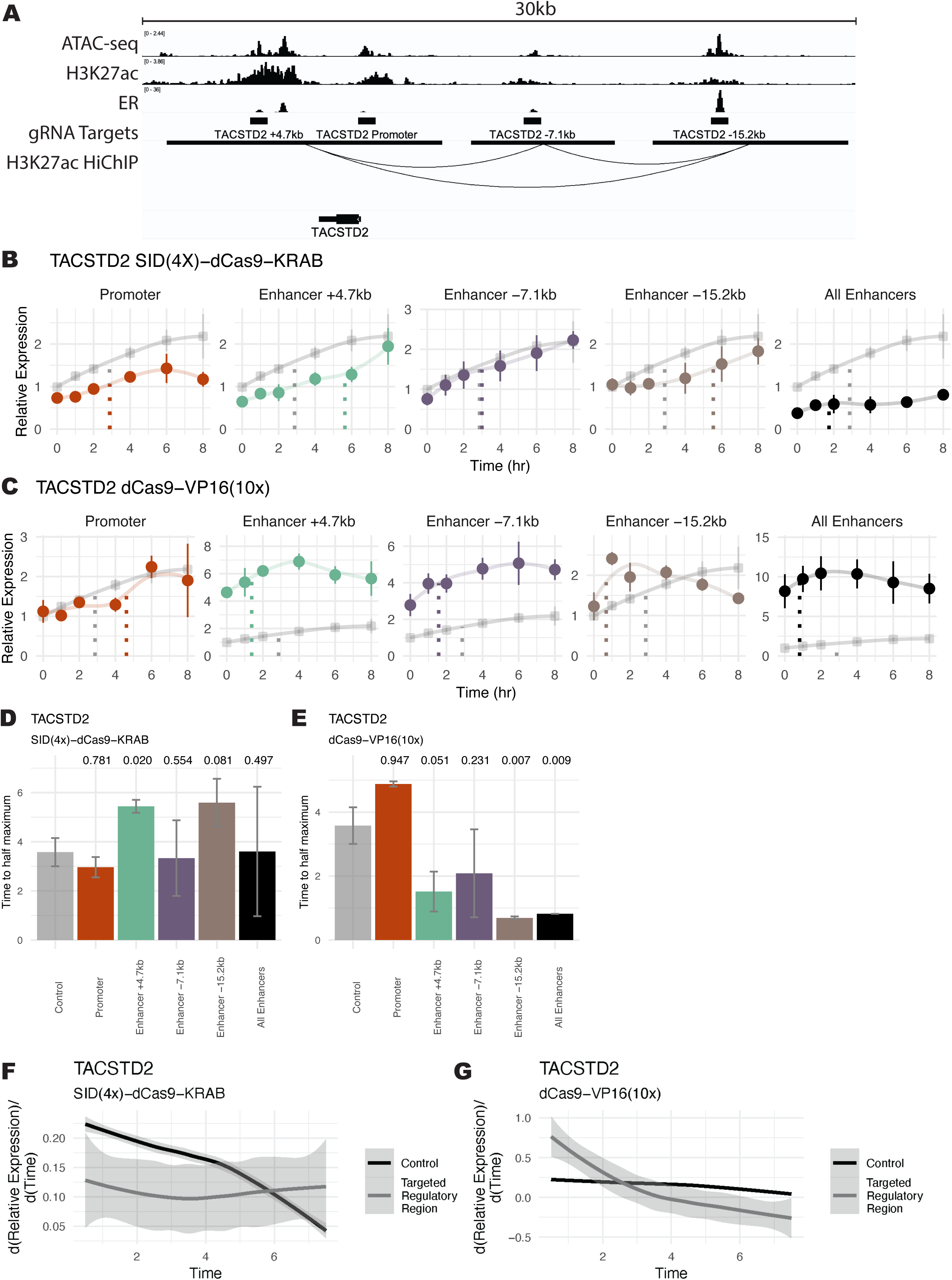
Functional manipulation of enhancer activity alters *TACSTD2* E2 response timing. (A) ChIP-seq, ATAC-seq, and HiChIP genome browser tracks in Ishikawa cells show targeted regulatory regions surrounding *TACSTD2*. (B-C) Expression trajectory of *TACSTD2* is displayed after E2 induction in Ishikawa cells following SID(4x)-dCas9-KRAB inhibition (B) or dCas9-VP16(10x) activation (C) targeted to regulatory regions. Error bars represent SEM (n=2) and expression is relative to the 0-hour timepoint in the control, which is from cells with *IL1RN* promoter targeting. Dotted lines show the time at half maximum for a given trajectory. (D-E) Bar plot shows time to half maximal expression for each targeted regulatory region. Error bars represent SEM and p-values (one-sided t-test) are reported above each bar. (F-G) Aggregate differential of loess regressions from B and C for all regulatory regions targeted (grey) by SID(4x)-dCas9-KRAB (F) or dCas9-VP16(10x) (G) compared to control (black). Shaded region represents 95% confidence interval.

When targeting five putative enhancers as well as the promoter of *TGFA*, an Early Up gene, with SID(4X)-dCas9-KRAB, we again observed a more gradual expression response to E2 (Figure S4A). The most substantial effects on time to half maximal expression were observed when targeting enhancer −43kb, enhancer −62kb, or all enhancers simultaneously (Figure S4C). On aggregate, inhibition of *TGFA* regulatory regions showed slower activation rates between early timepoints, followed by increased rates from 6 to 8 hours relative to the control trajectory (Figure S4E). These results are consistent with our *TACSTD2* findings and indicate that decreasing enhancer activity slows the transcriptional response. To speed up a Late gene, dCas9-VP16(10x) was used to activate enhancers and the promoter of *PEG10*. We observed an overall faster response when targeting these regulatory elements, although baseline levels were increased in all cases, which resulted in more variability in the timing analysis (Figure S4B and D). On aggregate, we see that activation of *PEG10* enhancers led to earlier responses to E2, which later converge with control rates (Figure S4F). Overall, at these three genes, the activity of enhancers controls the E2 response trajectory.

### An enhancer-promoter dichotomy controls gene expression noise

Genes were separated into three levels of variation to find determinants of expression noise at the 0-hour timepoint. In scRNA-seq, low gene expression levels often have high noise due to the dropout effects of capturing RNAs from the limited amount of RNA in a single cell and technical variation in scRNA-seq is related to mean levels (Brennecke et al., 2013). To examine mean-independent noise, we used an adjusted coefficient of variation (CV), which is calculated as the residuals of a generalized additive model (GAM) where CV is fit to the mean (Figure S5A, see Methods). To remove any leftover mean effects, genes were labeled as high or low noise based on whether they were in the top 20% or bottom 20% of adjusted CV for ten different mean bins from the 0-hour timepoint of both cell types (Figure S5B).

The strongest predictors of noise levels were SIN3A and JUN at the promoter, both associated with low noise (Figure 4A, B, and D). Generally, a strong promoter signal was related to low noise across features, with some exceptions, such as p300 (Figure 4A, right panel). Most enhancer features were associated with high noise, with ER and FOXA1 at enhancers being the most predictive (Figure 4A). Another feature scored as highly important was tri-methylation at histone H3 lysine 4 (H3K4me3), which is strongly associated with low amounts of noise (Figure 4C), supporting a previously reported correlation between H3K4me3 breadth and transcriptional consistency (Benayoun et al., 2014). The above analysis was performed on the combined Ishikawa and T-47D data; however, the importance scores of features when run separately on the cell types were correlated (r = 0.58; Figure S5C), indicating that common features control transcriptional noise.

**Figure 4.**
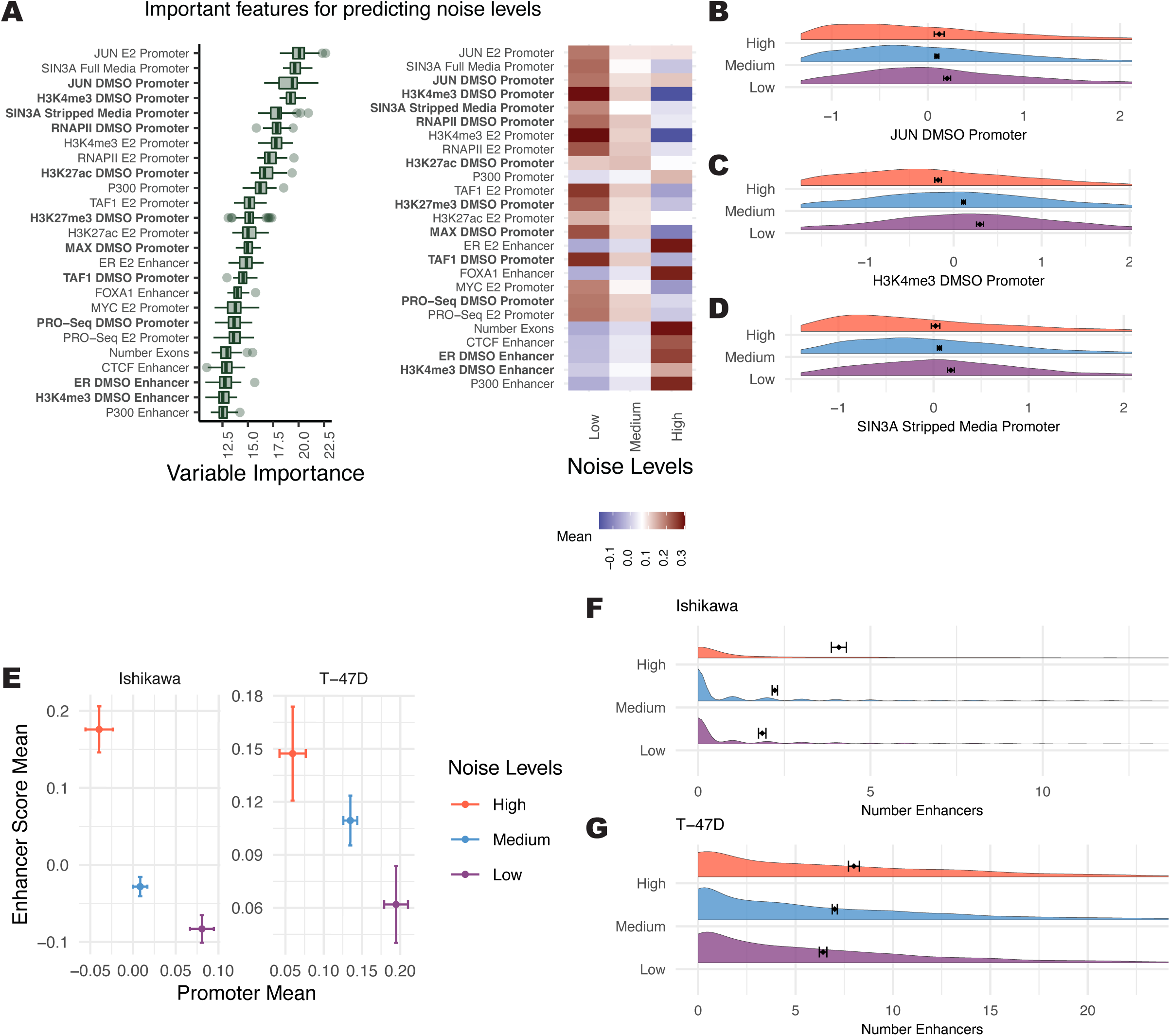
Features associated with noise levels indicate a balance between active promoters and active enhancers. (A, left) Boruta feature rankings shows features predictive of noise levels at the 0-hour timepoint. (A, right) Average signal intensity is shown by noise group for top ranked features. Datasets shown in bold were performed in the absence of ER activation. (B-D) Distribution of signal for top ranked noise-predicting features in the absence of ER activation are shown with Z-scores on the x-axis. (E) Mean enhancer signal score for all Boruta confirmed features vs. mean promoter signal across all confirmed features for each noise level exhibits an inverse relationship. Error bars show 95% confidence intervals and axes do not represent the full range; full distribution shown in Figure S6A. (F-G) Distribution of enhancer counts per gene, separated by noise level, are shown for (F) Ishikawa and (G) T-47D cells.

These results motivated the broader evaluation of how noise relates to promoter and enhancer activity. Analysis of the average promoter intensity across all confirmed datasets and the average enhancer score revealed an inverse relationship between promoters and enhancers (Figures 4E and S6A). Genes with high noise levels had high enhancer scores and low promoter signals. Conversely, genes with low noise levels had low enhancer scores and high promoter signals. To confirm this relationship in a third cell type, we analyzed publicly available scRNA-seq and genomic data from LNCaP cells, a prostate cancer cell line. The same association was observed between enhancer-driven gene regulation and higher noise (Figure S5D). Consistent with these findings, more enhancers connected to a gene associates with greater noise (Figure 4F and G). These results indicate that enhancer-driven transcription regulation is less consistent across individual cells than promoter-driven transcription regulation.

### Shared features of expression levels, timing, and noise

Expression levels, noise, and timing analysis uncovered different importance rankings for genomic features. Importance scores fell roughly into five common patterns across our three analysis types (Figure 5A). The largest three patterns consisted of features whose importance scores were highly enriched for a single analysis. Two smaller patterns were composed of features important for both mean and noise or both trajectory and noise. In general, the importance scores from the Boruta algorithm are more similar between noise and mean or noise and trajectory compared to mean and trajectory, as seen from the first two PCA dimensions calculated from the feature importance matrix (Figure 5B) and the correlation between importance scores (Figure S6D-F). The relationship between noise and our other analyses suggests that noise may be an intermediary between baseline levels and temporal regulation and that mean levels do not strongly influence response trajectory. Generally, we see that most features specifically associated with mean levels are found at promoters, while noise and trajectory utilize promoter and enhancer features more evenly (Figure 5A). Enhancer features important for predicting mean levels are also likely to predict noise. Together, these results indicate that enhancers are more critical for noise and trajectory and that many genomic signals preferentially predict either levels, noise, or trajectory.

**Figure 5.**
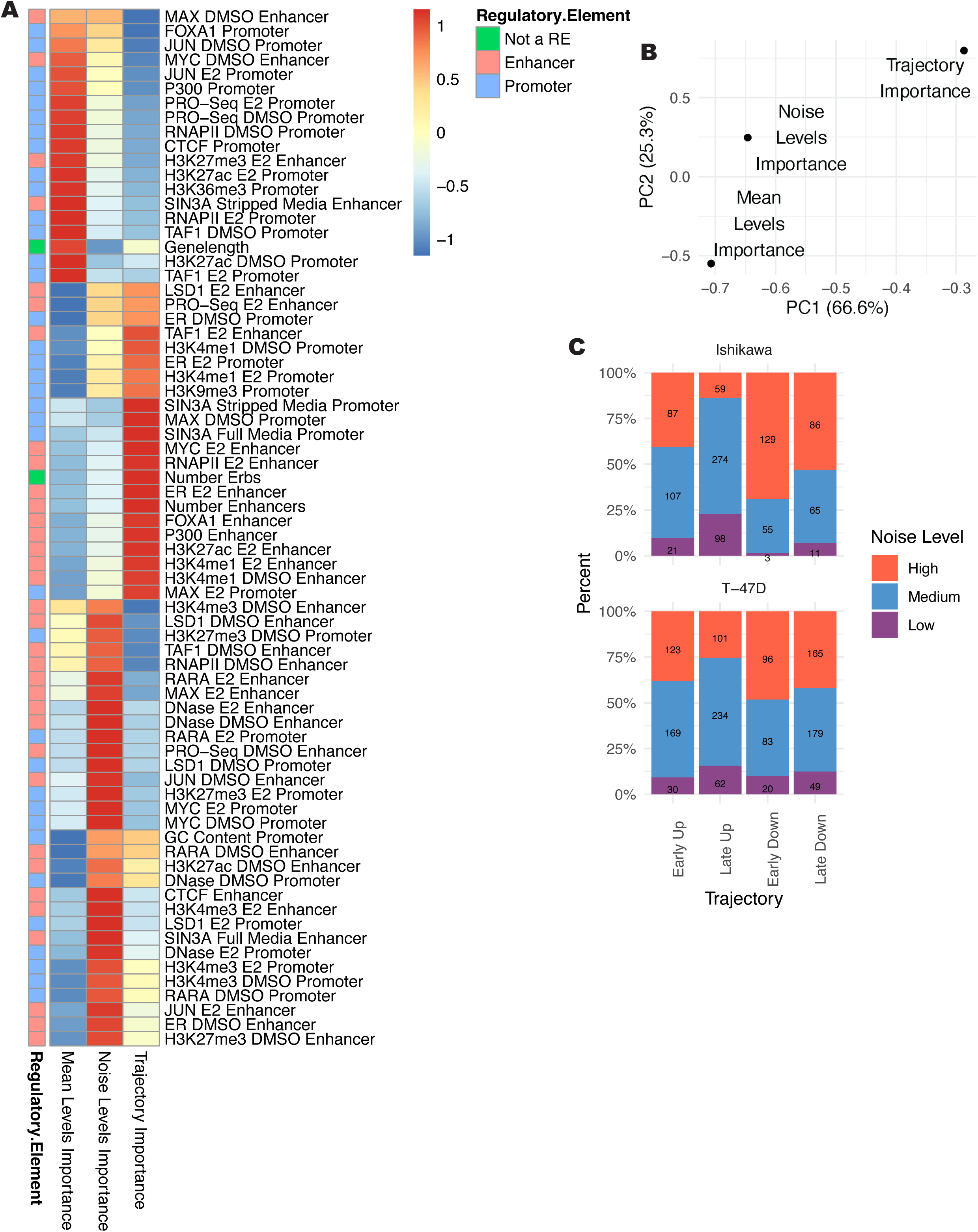
Importance comparison shows that mean and trajectory are regulated by distinct genomic features. (A) Heatmap shows importance scores from each analysis type, normalized by column, and scaled by row. Datasets shown in bold were performed in the absence of ER activation. (B) PCA plot based on importance scores shows the relationship of importance scores for mean levels, noise, and trajectory. Percentages denote percent of variance explained by each principal component. (C) Bar plot shows the percentage of genes for each trajectory that are classified into each noise classification. Numbers on bars refer to counts of genes.

Since Boruta importance scores do not capture directionality, we examined the group with the maximum signal for each feature (Figure S6C). Promoter features are generally associated with high mean levels and low noise. Enhancer features are also associated with increased mean levels, but contrary to promoters, they show an association with high noise levels. Enhancer scores are almost always the highest for Early Up trajectories. As high enhancer scores are associated with Early Up trajectories and high noise, we performed a gene level comparison of noise at the 0-hour timepoint and trajectory classification. We found that genes that respond quickly to E2 treatment are more likely to exhibit high noise at the 0-hour timepoint (Figure 5C). Our results suggest that active enhancers drive high noise and rapid up-regulation in response to E2, while promoters consistently drive low noise and high mean expression.

### Co-expression of genes is associated with looping, timing, and noise levels

scRNA-seq offers a unique advantage in studying the co-regulation of genes on a cell-by-cell basis and the possible mechanisms that underlie co-regulation. Using the H3K27ac HiChIP data, we found that looping can affect co-expression in several ways. First, we found that genes whose promoters loop together correlate significantly more than groups of randomly paired control genes at the 0-hour timepoint (Figure 6A and B). Genes whose promoters both loop to a shared enhancer are significantly more correlated across single cells (Figure 6C and D). These results indicate that the 3D genome structure may be involved or associated with gene co-expression across single cells.

**Figure 6.**
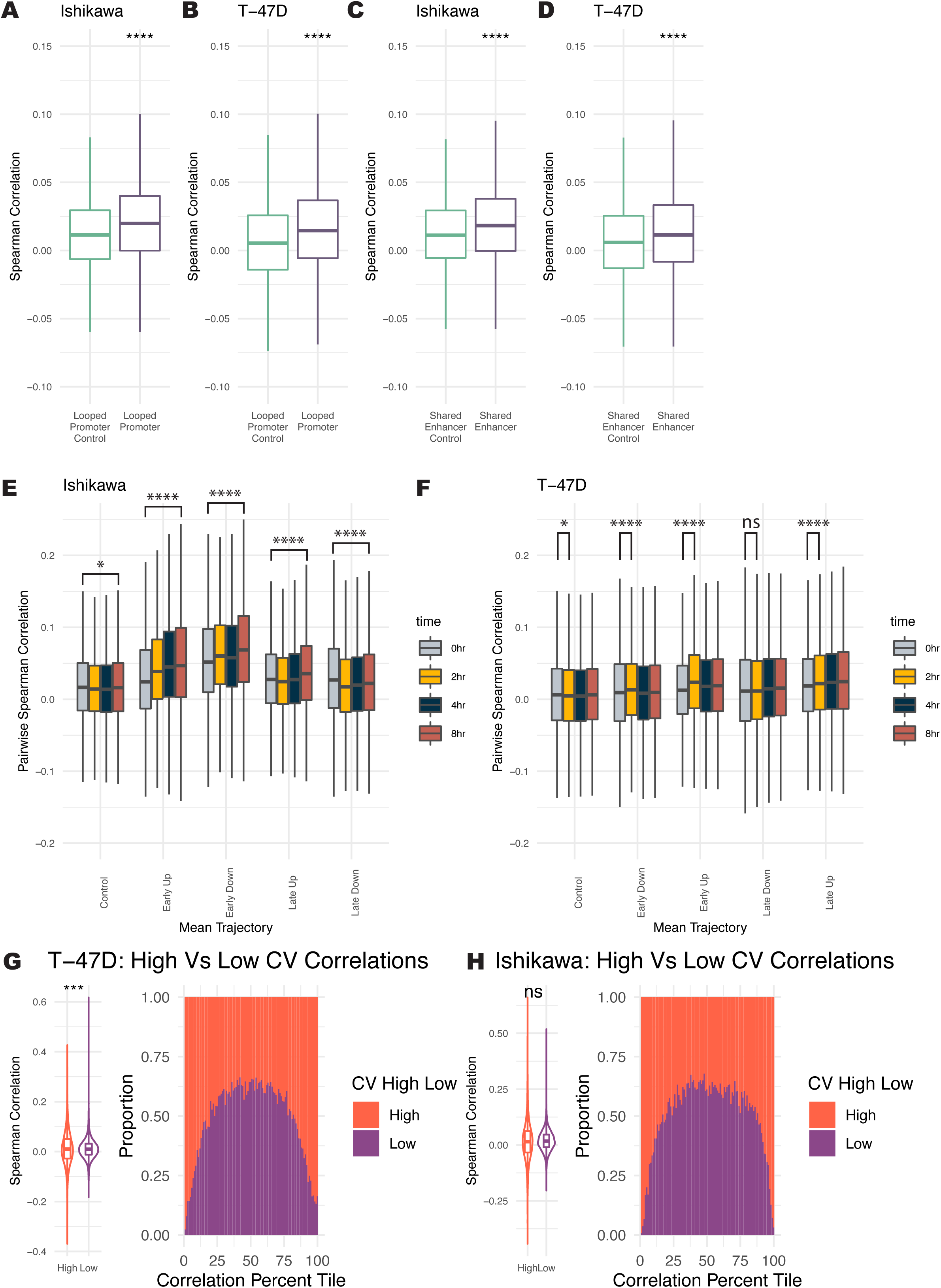
Co-expression levels track with looping, trajectory, and levels of noise. (A-B) Pairs of genes with promoters that loop to one another are significantly more correlated across cells at the 0-hour timepoint than randomly selected gene pairs for Ishikawa (A) and T-47D (B). (C-D) Pairs of genes with a shared enhancer are more correlated than randomly paired genes for Ishikawa (C) and T-47D (D). (E-F) Distribution of pairwise Spearman correlation for genes within different trajectories is shown for Ishikawa (E) and T-47D (F). (G-H) Range of pairwise correlations for high noise levels is greater than the range for pairs of low noise genes in Ishikawa (G) and T-47D (H). (left panel) Distribution of Spearman pairwise correlations for genes with high and low noise. (right panel) Spearman correlations were grouped into quantiles and bars show proportion at each quantile that are pairs of low or high noise genes. Significance values for all subpanels are as follows (based on Bonferroni corrected Wilcoxon tests): (* p < 0.05; ** p < 1×10-5; *** p < 1×10−10; **** p < 1×10−15).

We next evaluated co-expression during the E2 treatment time course. Co-expression was measured using pairwise Spearman correlation in single cells. In Ishikawa cells, both Early Up and Early Down genes show increasing pairwise co-expression over time (Figure 6E). Genes that respond late exhibited less change in correlation, with Late Up genes increasing correlation slightly by 8 hours and Late Down genes slightly decreasing in correlation. In T-47D cells, we see the most significant increase in co-correlation at 2 hours for Early Up and Early Down genes (Figure 6F). Late Up and Late Down genes show slight increases in correlation during the time course.

The levels of noise also change the probability of two genes being correlated. Perhaps expected, genes with high noise levels also show a broader distribution in their pairwise correlations, resulting in genes with high noise being more likely to have extremely correlated or anti-correlated expression with each other than low noise genes (Figure 6G and H). These results indicate that 3D interactions, control of temporal trajectory, and noise regulation can impact gene co-regulation.

## Discussion

To investigate the genomic underpinnings of the temporal transcriptional response to estrogen, we conducted scRNA-seq at several timepoints in two human cell lines. scRNA-seq was able to capture more subtle changes in gene expression of highly expressed genes compared to bulk RNA-seq due to the increased statistical power. Using a feature ranking approach, we identified several features associated with E2 response timing, including more ER bound enhancers regulating “Early Up” genes and SIN3A signal at the promoter of “Late Up” genes. In general, multiple enhancers are more predictive of “Early” gene trajectories. Functional evaluation of enhancers revealed that multiple enhancers regulate timing at each gene tested as activation and repression of several enhancers causes changes in a gene’s temporal response to estrogen. From these studies, we conclude that an active enhancer repertoire is important for rapid gene responses to estrogen. Active enhancers may present chromatin that is more open to ER binding. Alternatively, other TFs already present at enhancers could stabilize the binding of ER, permitting ER to activate gene expression immediately. Contrastingly, SIN3A and MAX at the promoter, known to repress gene expression together (Baudino and Cleveland, 2001), slow a gene’s response to E2, even when a gene is associated with strong ER bound enhancers. Activation of gene expression may first require removing repressive signals at the promoter, explaining the more gradual responses. A similar mechanism has been described in an enhancer context, where inactive enhancers must first be activated by transcription factors before activation of gene expression can occur, causing more gradual gene expression changes (Ostuni et al., 2013). While transcriptional responses to estrogen (Gertz et al., 2012) as well as underlying gene regulatory events (Gertz et al., 2013) are highly cell type-specific, we found that the genomic features associated with the transcriptional response are shared between cell types, indicating that common mechanisms drive how genes respond to estrogen stimulation.

It is important to note that we analyzed total poly-adenylated RNA in this study. Since mRNA degradation can happen on different time scales across genes (Tani et al., 2012), some genes may be misclassified in their transcriptional response to estrogen. In fact, we found that Early Down genes had shorter mRNA half-lives in previously published datasets. This observation is consistent with quickly down-regulated genes requiring fast mRNA turnover to be observed from poly-adenylated RNA, since overall transcript levels would need to be significantly reduced in a short time period. The confounding effect of mRNA stability on down-regulated genes may partially explain why we did not find features that consistently associate with Early/Late Down genes. The lack of strong association between the genomic features we studied and down-regulated genes could also be related to a focus on features related to gene activation and a stronger connection between ER and gene activation as opposed to repression (Carleton et al., 2017).

One unexpected and exciting finding from our analysis is that enhancer and promoter activities relate to expression noise in opposing manners. Active promoters are associated with low noise levels, whereas multiple active enhancers are associated with high noise levels. In concordance with this observation, synthetic activation of promoters drives lower noise levels at several genes (Fraser et al., 2021a). Additionally, activation of multiple enhancers causes high noise at the NF-κB locus (Wibisana et al., 2022). Our results support a unified model where a balance between enhancers and promoters controls noise. Both intrinsic and extrinsic noise could potentially explain the observed noise distributions (Ham et al., 2021). If intrinsic noise is the driving factor, we expect promoters to cause high-frequency, near-constant transcription levels and enhancers to cause infrequent, high-amplitude bursts of expression (Raj and Van Oudenaarden, 2008). Noise caused by CREs could also be due to extrinsic noise. Promoters may lead to low noise, as fluctuations in upstream factors may be insignificant compared to activation by an ensemble of transcription factors bound to the promoter. In contrast, enhancers may drive higher noise levels by increasing sensitivity to upstream factors, which is consistent with our observation that enhancers drive rapid temporal responses to estrogen.

For a gene to respond quickly to a signal, it must be sensitized to incoming signals, which may inherently drive higher levels of noise. A noise-robustness tradeoff has been proposed previously when observing changes in gene expression over developmental time in *drosophila* (Sigalova et al., 2020) and in a mathematical framework that showed variation is necessary for a gene’s responsiveness (Boe et al., 2022). However, a regulatory mechanism has not been found. Our results point to multiple enhancers being a primary genomic feature associated with both high expression noise and rapid response timing. Additionally, we found that a strong promoter is likely to cause more robust gene expression but limited responsiveness. While we identified these patterns across the full data set, a small set of individual genes can respond quickly to a signal without high cell-cell variation (Figure 5C) and the question remains as to how these genes maintain low noise and fast response times. We also found that genes that are quickly down-regulated by estrogen are more likely to exhibit high noise than genes that are quickly up-regulated by estrogen. This pattern could be due to genes that are primarily enhancer driven having higher noise and being more susceptible to down regulation, potentially through enhancer rewiring because of ER activity and competition for factors that are critical for enhancer activity. Shorter mRNA half-lives could also contribute to higher noise of Early Down genes.

Co-expression analysis of gene pairs showed that co-expression properties depend on looping, timing, and noise. We found that genes with shared enhancers and looped promoters correlate more in individual cells, genes with different trajectories correlate differently over time, and genes with high noise levels are more likely to be strongly correlated or anticorrelated. While co-expression correlation effects are modest overall and it is unclear what levels of co-expression are biologically meaningful, these observations could have implications for gene regulatory networks (GRNs) in single cells, as co-expression often underlies regulatory networks. Dynamic adjustment of regulatory networks may have critical functional outcomes for a cell population (Borriello et al., 2020). For example, GRNs that confer resistance to therapeutics may occur at distinct timepoints following treatment (Zhang et al., 2019). Our results indicate that genes with high noise may lead to a broader range of implemented regulatory networks across single cells, enhancing cellular heterogeneity. Further studies into functional GRNs are warranted to determine how noise and timing affect single cell phenotypes through the co-expression of many genes. Overall, our study shows that enhancers and promoters can play distinct roles in the timing and variation of a transcriptional response.

## Methods

### Cell culture

T-47D and Ishikawa cells were cultured in RPMI 1640 medium (Gibco) with 10% fetal bovine serum (Gibco) and 1% penicillin–streptomycin (Gibco). LNCaP cells were cultured in RPMI media with 10% FBS supplemented. Cells were incubated at 37°C with 5% CO_2_. 5 days before estrogen inductions, cells were transferred to hormone-depleted media consisting of phenol red-free RPMI (Gibco) with 10% charcoal-dextran stripped fetal bovine serum (Sigma-Aldrich) and 1% penicillin–streptomycin (Gibco).

### ChIP-seq

After 5 days in hormone-depleted media, cells were plated in 15cm dishes at approximately 60% confluency 1 day before estrogen induction. Cells were treated with vehicle (DMSO) or E2 at a final concentration of 10nM for either 1 hour for transcription factor ChIP-seqs, or 8 hours for histone marker ChIP-seqs. ChIP and library preparation was performed as previously described (Reddy et al., 2009). Antibodies used for this study were MAX (Sant Cruz sc-8011), LSD1 (abcam ab 17721), TAF1 (sc-735), c-MYC (Santa Cruz sc-40), H3K4me3 (Cell Signaling 9751S), H3K4me1 (Cell Signaling 5326S), SIN3A (produced as previously described) (Hassig et al., 1997), RARA (Santa Cruz sc-515796) and JUN (BD Biosciences 558036). Libraries were sequenced using either an Illumina HiSeq 2500 or Illumina NovaSeq 6000 as single-or paired-end 50 bp reads, then aligned to hg19 using bowtie with parameters -m 1 –t – best -q -S -l 32 -e 80 -n 2 (Langmead et al., 2009). Signal intensity was extracted from bam files using samtools view with parameter -c (Li et al., 2009). In the cases where peaks were called, peak calling was done using Macs2 with the default q-value cutoff of 0.05 and mfold ratio between 15 and 100 (Zhang et al., 2008).

### H3K27ac HiChIP

HiChIP experiments were performed as previously described (Mumbach et al., 2016) using an antibody that recognizes H3K27ac (Abcam, ab4729). Ishikawa cells were treated with either 10 nM E2 for 1 hour or DMSO as a vehicle control. HiChIP in Ishikawa cells was conducted using restriction enzyme DpnII (New England Biolabs). Crosslinked chromatin was sonicated using an EpiShear probe-in sonicator (Active Motif) with three cycles of 30 seconds at an amplitude of 40% with 30 seconds rest between cycles. HiChIP libraries were sequenced on NovaSeq 6000 as paired end 50 base pair reads to an average depth of 300–400 million read-pairs per sample.

Experiments in T-47D and LNCaP cells were conducted using restriction enzyme MboI (New England Biolabs). Crosslinked chromatin was sonicated using Covaris E220 with the settings of fill level=10, duty cycle=5, PIP=140, cycles per burst=200, time=4 mins. HiChIP libraries were sequenced on HiSeq 2500 as paired end 75 base pair reads to ∼50 million read pairs per sample.

Reads were aligned to human hg19 reference genome using HiC-Pro (Servant et al., 2015). Hichipper (Lareau and Aryee, 2018) was used to perform restriction site bias-aware modeling of the output from HiC-Pro and to call interaction loops. In Ishikawa cells, DMSO and E2 treated HiChIP loops were combined to identify all possible putative enhancers. In all datasets, loops with less than 3 reads or FDR >= .05 were filtered out.

### PRO-seq

PRO-seq libraries were generated as described in Mahat et al., 2016 (Mahat et al., 2016). Briefly, Ishikawa and T-47D cells were grown in hormone-depleted RPMI for five days, then 2×10^6^ cells were plated into two 10 cm dishes per condition with RPMI lacking phenol red supplemented with 10% charcoal/dextran-stripped FBS and penicillin. Cells were treated with vehicle (DMSO) or 10 nM E2 for 45 minutes, then permeabilized for five minutes with permeabilization buffer [10 mM Tris-HCl, pH 7.4; 300 mM sucrose; 10 mM KCl; 5 mM MgCl2; 1 mM EGTA; 0.05% Tween-20; 0.1% NP40 substitute; 0.5 mM DTT, protease inhibitor cocktail ml(Roche); and SUPERaseIn RNase Inhibitor (Ambion)]. The nuclear run-on was performed by adding permeabilized cells to run-on mixture [final composition was 5 mM Tris, pH 8.0; 25 mM MgCl2; 0.5 mM DTT; 150 mM KCl; 200 μM rATP; 200 μM rGTP; 20 μM biotin-11-rCTP (Perkins Elmer); 20 μM biotin-11-rUTP (Perkins Elmer); 1 U/μL SUPERase In RNase Inhibitor (Ambion); 0.5% Sarkosyl], then incubating at 37°C for 5 minutes. RNA was extracted with Trizol LS (Ambion), fragmented with 0.2 N NaOH for 8 minutes on ice, then neutralized with 0.5 M Tris, pH 6.8, followed by buffer exchange with a P-30 column (Bio-Rad). Biotinylated RNAs were enriched with Dynabeads M280 Streptavidin (Invitrogen), then RNA was extracted with Trizol (Ambion), followed by 3′ adapter ligation using T4 RNA ligase (NEB). Biotinylated RNAs were enriched for a second time, followed by 5′ cap repair with RppH (NEB) and 5′ hydroxyl repair with PNK (NEB). The 5′ adapter was ligated with T4 RNA ligase (NEB), followed by a third biotinylated RNA enrichment. Reverse transcription was performed with the RP1 primer. Samples were PCR amplified for 13 cycles, then cleaned up with Agencourt AMPure XP beads (Beckman Coulter). Libraries were sequenced on an Illumina HiSeq 2500, generating a 50nt read. Reads were processed using cutadapt (Martin, 2011) with parameters -a TGGAATTCTCGGGTGCCAAGG --cut 7 --length 42 -m 21. Reverse complement sequences were generated using fastx_reverse_complement from the FASTX toolkit (v 0.0.13) (Hannon, 2010). Reads were then aligned to hg19 with bowtie2 (Langmead and Salzberg, 2012) in end-to-end mode, and non-uniquely aligned reads were discarded.

### scRNA-seq

Cells were treated with 10nM E2 for 0 (vehicle treated), 2, 4, and 8 hours. To mitigate technical batch effects, cells were labeled via MULTI-seq as previously described (McGinnis et al., 2019). Cells from different timepoints were mixed and then prepared according to the 10x Genomics sample prep user guide (2017). Cells were separated into single cell emulsions using the 10x Genomics Chromium Controller with a targeted recovery of 10,000 cells. Sequencing libraries were prepared using the 10X Genomics Next GEM Single Cell 3’ Gene Expression Library prep v3.1. Sequencing was performed on an Illumina NovaSeq 6000 with 150bp read length. Sequencing output was processed from reads to counts using the 10x Genomics Cell Ranger v3.1.0 pipeline. MULTI-seq calls were processed using the demultiplex R package (McGinnis et al., 2019) and mapped back to the E2 timepoints. Counts were log normalized using the Seurat v3 R package (Stuart et al., 2019), then filtered using custom cutoffs. For Ishikawa cells, cell filtering criteria were unique reads between 8500 and 35,000, unique genes between 2700 and 6000, and percent mitochondrial reads less than 7%. For T-47D cells, cell filtering criteria were unique reads between 7500 and 40,000, unique genes between 1500 and 6000, and percent mitochondrial reads less than 20%. Genes are filtered to have a mean greater than 0.01 across all timepoints.

### scRNA-seq analysis: classification of trajectory and noise levels

Computational analysis of trajectory and noise levels were conducted using R (Team, 2013). Trajectory classification was done using a Wilcoxon test (Bauer, 1972) to find genes whose single cell distributions significantly change at different timepoints compared to the 0hr timepoint. Genes that change significantly by 2 hours are classified as either “Early Up” or “Early Down”. Genes with changes seen at 4 or 8 hours are classified as “Late”. It is important to note that genes were called differential without a fold change cutoff. The statistical power of scRNA-seq allowed for the identification of differentially expressed genes with smaller fold changes, but appreciable absolute changes; however, due to technical limitations of scRNA-seq our results may be affected by technical variation or drop out of lower expressed genes. To select control genes with similar mean distributions to those genes that are regulated by E2, we used a stratified sampling approach to select control genes that are not significantly regulated. Genes classified into Early and Late Up categories were analyzed by EnrichR (Xie et al., 2021) using the MSigDB Hallmark 2020 gene annotations.

Our noise metric is defined as the residuals from a generalized additive model (GAM) regression fitted to the CV vs mean for all genes. Regression was performed on log2(CV + 1) vs mean curve using the gam function from the mgcv R package with formula y ∼ s(x, bs = “cs”) (Wood, 2006). Residuals were then transformed back to the original scale. Noise levels were determined using the GAM-adjusted CV. We chose the GAM method as it removed most of the mean-noise relationship (Figure S6B), as desired, compared to SCTransform standardized variance and residual variance as shown in Figure S5E and F. Overall, we found that genes were binned as High noise more consistently across the three methods (average fraction overlap of 0.82) than Medium (0.67) or Low noise (0.58); however, less than 1% of genes were classified as High noise by one method and Low noise by another. For this comparison of noise metrics, we used the HVFInfo function with selection.method as “vst” or “sctransform” in Seurat (Satija et al., 2015). To account for the different ranges in noise at different mean levels, genes were binned by mean and then into 3 groups of noise levels by quantile. The top 20% and bottom 20% of genes in each quantile were labeled as “High” and “Low” noise, respectively.

### Feature importance analysis

Promoters were defined as 500bp regions centered on the transcriptional start site, as annotated in the RefSeq database (Pruitt et al., 2013). Enhancers were called using H3K27ac HiChIP data and H3K27ac ChIP-seq peaks. Enhancers were defined as H3K27ac peaks within HiChIP anchors that loop to the promoter. Integrated signal for each promoter and enhancer was collected from all datasets using samtools view -c (Li et al., 2009). Z-scores were calculated across all genes for input to feature ranking algorithms. An enhancer score was calculated to account for signal at multiple enhancers, using the formula

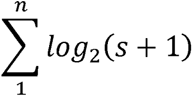

where n represents the number of enhancers associated with a gene and s represents the Z-score of integrated genomic signal at each enhancer.

Number of enhancers was defined as the number of H3K27ac peaks that loop to the promoter, as determined by HiChIP. Number of ERBS was calculated as the number of enhancers that overlapped with ER ChIP-seq peaks. Feature ranking was performed using the Boruta package in R (Kursa and Rudnicki, 2010b) with default parameters and 100 maximum iterations. Gene length was calculated from RefSeq transcript annotations (Pruitt et al., 2013). An example decision tree was determined using the rpart function with parameters minbucket=50 and cp=0.007 (Breiman et al., 2017). Average enhancer score and promoter signal was calculated using “confirmed” variables from Boruta analysis. The average of Z-score signal for confirmed variables was taken for all variables associated with either the promoter or enhancer, not including number of enhancers, number of ERBS, or gene length.

### Generation of stable dCas9-VP16(10x) cell lines

Ishikawa cells were plated in 6-well plates at 60% confluency. Cells were transfected with Addgene plasmid 48227 (a gift from Rudolf Jaenisch) (Cheng et al., 2013) containing dCas9-VP16(10x) with a P2A linker and neomycin resistance gene. Fugene HD (Promega) was used for transfection at a 3:1 reagent:DNA ratio. dCas9-VP16(10x) plasmid was linearized with restriction enzyme AflII (New England Biolabs R0520S). Successful integration of the dCas9-VP16(10x) plasmid was selected for using G418 (Thermo Scientific) at a concentration of 800 µg/mL for approximately 2 weeks. Successful expression of the dCas9 plasmid was verified using qPCR for dCas9 as well as qPCR for successful activation of a control gene, *IL1RN* (Figure S4G). Cells were then maintained at a lower concentration of 400 µg/ml G418.

### gRNA design and transfection

gRNAs were designed using the Benchling gRNA design tool (Benchling, 2020). 4 gRNAs were designed per targeted region. gRNAs were cloned into plasmids as previously described (Carleton et al., 2017). gRNA sequence and adjacent PAM are listed in Table S1. Prior to transfection, Ishikawa cells were plated in 48-well plates at 80,000 cells/well. 24 hours after plating, gRNAs were transfected into cell using Fugene HD (Promega) at a manufacturer suggested 3:1 reagent:DNA ratio. gRNA transfection was selected for using 1 µg/mL puromycin. 8-hour E2 time courses were started roughly 24 hours after addition of puromycin.

### RNA isolation and qPCR gene expression analysis

Cells were lysed with Buffer RLT Plus (Qiagen) containing 1% beta-mercaptoethanol (Sigma). RNA was purified using the ZR-96-well Quick-RNA kit (Zymo Research). Gene expression was measured using qPCR with reagents from the Power SYBR Green RNA-to-Ct 1-step kit (Applied Biosystems), 50ng RNA per reaction, and 40 cycles on a CFX Connect light cycler (BioRad). qPCR primers are listed in Table S2. Relative expression was calculated using the ΔΔCt method with *CTCF* as a reference gene and cells where the same dCas9 fusion is targeted to the *IL1RN* promoter as the controls. Best fit lines were determined using the loess function in R (Cleveland et al., 2017) and formula y ∼ x. Half maximal values were calculated using the maximum at any timepoint. The first time point at which the loess regression reaches a half maximal value is recorded as the time to half maximal. Differentials of the loess regression were also calculated using R to examine how the slope of each trajectory changes over time. Slope analysis was binned into “targeted” and “control” groups to analyze the general effect of dCas9 manipulation at regulatory regions of the respective gene.

### Bulk RNA-seq experiments and analysis

Cells were treated with 10nM E2 for 0 (vehicle treated), 2, 4, and 8 hours. Following treatments, cells were lysed with buffer RLT Plus (Qiagen) containing 1% beta-mercaptoethanol (Sigma-Aldrich). RNA was extracted and purified using a Quick RNA Mini Prep kit (Zymo Research). NEBNext Ultra II Directional RNA Library Prep kit with poly(A) mRNA isolation was used to construct RNA-seq libraries according to the manufacturer’s instructions (NEB). Sequencing reads were aligned to hg19 build of the human genome using HISAT2 (Kim et al., 2019). SAMtools (Li et al., 2009) was used to convert SAM files to BAM files. Genes were defined by the University of California Santa Cruz (UCSC) Known Genes (Kent et al., 2002) and reads that mapped to known genes were assigned with featureCounts (Liao et al., 2014). Read counts were normalized and analyzed for differential expression via DESeq2 (Love et al., 2014). Genes that were differentially expressed between the 0-hour and 2-hour timepoints were defined as Early, while all other differentially expressed genes were defined as Late.

### Co-expression analysis

Spearman correlation values were calculated using the cor function in R. Pairs were defined either by looping data, or for a set of genes, as all possible combinations of genes in a list. Control pairings for genes that were looped together were generated by randomly shuffling the pairings. Control genes in the time course correlation analysis refer to the pairs within the set of timing control genes (defined above) where stratified sampling was used to replicate the same mean distribution as regulated genes.

## Author Contributions

Conceptualization, M.G-H. and J.G.; Methodology, M.G-H. and J.G.; Investigation, M.G-H., H.A., K.L.M., and E.M.W; Formal Analysis, M.G-H., J.G., and X.Z.; Writing - Original Draft, M.G-H. and J.G.; Writing – Review & Editing, All authors; Supervision, J.G., J.L., and X.Z.; Funding Acquisition, J.G.

## Acknowledgements

Funding for this work came from National Institute of Health (NIH)/National Human Genome Research Institute (NHGRI) R01 HG008974 to J.G. and the Huntsman Cancer Institute. Research reported in this publication utilized the High-Throughput Genomics Shared Resource at the University of Utah and was supported by NIH/National Cancer Institute (NCI) award P30 CA042014. We thank Craig M. Rush for experimental guidance, Jeffery Vahrenkamp for analysis advice, and Gertz lab members for their suggestions on the study and manuscript.

## Data Availability

ChIP-seq, HiChIP, PRO-seq, RNA-seq, and scRNA-seq data are available at the Gene Expression Omnibus under accession GSE227245.

## References

(2017). Single Cell Suspensions from Cultured Cell Lines for Single Cell RNA Sequencing. 10x Genomics Document Number CG00054 Rev B.

Agarwal, V., and Kelley, D.R. (2022). The genetic and biochemical determinants of mRNA degradation rates in mammals. Genome biology 23, 245.

Azpeitia, E., Balanzario, E.P., and Wagner, A. (2020). Signaling pathways have an inherent need for noise to acquire information. BMC Bioinformatics 21, 462.

Basma, H., Soto-Gutiérrez, A., Yannam, G.R., Liu, L., Ito, R., Yamamoto, T., Ellis, E., Carson, S.D., Sato, S., Chen, Y., et al. (2009). Differentiation and transplantation of human embryonic stem cell-derived hepatocytes. Gastroenterology 136, 990–999.

Baudino, T.A., and Cleveland, J.L. (2001). The Max network gone mad. Mol. Cell. Biol. 21, 691–702.

Bauer, D.F. (1972). Constructing confidence sets using rank statistics. Journal of the American Statistical Association 67, 687–690.

Behar, M., and Hoffmann, A. (2010). Understanding the temporal codes of intra-cellular signals. Curr. Opin. Genet. Dev. 20, 684–693.

Benayoun, B.A., Pollina, E.A., Ucar, D., Mahmoudi, S., Karra, K., Wong, E.D., Devarajan, K., Daugherty, A.C., Kundaje, A.B., Mancini, E., et al. (2014). H3K4me3 breadth is linked to cell identity and transcriptional consistency. Cell 158, 673–688.

Benchling (2020). Benchling.

Bignotti, E., Zanotti, L., Calza, S., Falchetti, M., Lonardi, S., Ravaggi, A., Romani, C., Todeschini, P., Bandiera, E., Tassi, R.A., et al. (2012). Trop-2 protein overexpression is an independent marker for predicting disease recurrence in endometrioid endometrial carcinoma. BMC Clinical Pathology 12, 22.

Bjornstrom, L., and Sjoberg, M. (2005). Mechanisms of estrogen receptor signaling: convergence of genomic and nongenomic actions on target genes. Mol. Endocrinol. 19, 833–842.

Boe, R.H., Ayyappan, V., Schuh, L., and Raj, A. (2022). Allelic correlation is a marker of trade-offs between barriers to transmission of expression variability and signal responsiveness in genetic networks. Cell Systems 13, 1016–1032. e1016.

Borriello, E., Walker, S.I., and Laubichler, M.D. (2020). Cell phenotypes as macrostates of the GRN dynamics. J. Exp. Zool. B Mol. Dev. Evol. 334, 213–224.

Breiman, L., Friedman, J.H., Olshen, R.A., and Stone, C.J. (2017). Classification and regression trees (Routledge).

Brennecke, P., Anders, S., Kim, J.K., Kołodziejczyk, A.A., Zhang, X., Proserpio, V., Baying, B., Benes, V., Teichmann, S.A., Marioni, J.C., et al. (2013). Accounting for technical noise in single-cell RNA-seq experiments. Nat. Methods 10, 1093–1095.

Carleton, J.B., Berrett, K.C., and Gertz, J. (2017). Multiplex Enhancer Interference Reveals Collaborative Control of Gene Regulation by Estrogen Receptor α-Bound Enhancers. Cell Syst 5, 333–344.e335.

Chamberlain, C.E., Jeong, J., Guo, C., Allen, B.L., and McMahon, A.P. (2008). Notochord-derived Shh concentrates in close association with the apically positioned basal body in neural target cells and forms a dynamic gradient during neural patterning. Development 135, 1097–1106.

Cheng, A.W., Wang, H., Yang, H., Shi, L., Katz, Y., Theunissen, T.W., Rangarajan, S., Shivalila, C.S., Dadon, D.B., and Jaenisch, R. (2013). Multiplexed activation of endogenous genes by CRISPR-on, an RNA-guided transcriptional activator system. Cell Res. 23, 1163–1171.

Choi, J.K., and Kim, Y.-J. (2009). Intrinsic variability of gene expression encoded in nucleosome positioning sequences. Nat. Genet. 41, 498–503.

Cleveland, W.S., Grosse, E., and Shyu, W.M. (2017). Local regression models. In Statistical models in S (Routledge), pp. 309–376.

Dadiani, M., Van Dijk, D., Segal, B., Field, Y., Ben-Artzi, G., Raveh-Sadka, T., Levo, M., Kaplow, I., Weinberger, A., and Segal, E. (2013). Two DNA-encoded strategies for increasing expression with opposing effects on promoter dynamics and transcriptional noise. Genome Res. 23, 966–976.

Desai, R.V., Chen, X., Martin, B., Chaturvedi, S., Hwang, D.W., Li, W., Yu, C., Ding, S., Thomson, M., Singer, R.H., et al. (2021). A DNA repair pathway can regulate transcriptional noise to promote cell fate transitions. Science 373, eabc6506.

Elowitz, M.B., Levine, A.J., Siggia, E.D., and Swain, P.S. (2002). Stochastic Gene Expression in a Single Cell. Science 297, 1183–1186.

Encode, C. (2012). An integrated encyclopedia of DNA elements in the human genome. Nature 489, 57–74.

Fidler, I.J. (1978). Tumor heterogeneity and the biology of cancer invasion and metastasis. Cancer Res. 38, 2651–2660.

Fraser, L.C., Dikdan, R.J., Dey, S., Singh, A., and Tyagi, S. (2021a). Reduction in gene expression noise by targeted increase in accessibility at gene loci. Proceedings of the National Academy of Sciences 118, e2018640118.

Fraser, L.C.R., Dikdan, R.J., Dey, S., Singh, A., and Tyagi, S. (2021b). Reduction in gene expression noise by targeted increase in accessibility at gene loci. Proceedings of the National Academy of Sciences 118, e2018640118.

Frasor, J., Danes, J.M., Komm, B., Chang, K.C.N., Lyttle, C.R., and Katzenellenbogen, B.S. (2003). Profiling of Estrogen Up-and Down-Regulated Gene Expression in Human Breast Cancer Cells: Insights into Gene Networks and Pathways Underlying Estrogenic Control of Proliferation and Cell Phenotype. Endocrinology 144, 4562–4574.

Fritzsch, C., Baumgärtner, S., Kuban, M., Steinshorn, D., Reid, G., and Legewie, S. (2018). EstrogenLdependent control and cellLtoLcell variability of transcriptional bursting. Mol. Syst. Biol. 14, e7678.

Gandhi, S.J., Zenklusen, D., Lionnet, T., and Singer, R.H. (2011). Transcription of functionally related constitutive genes is not coordinated. Nat. Struct. Mol. Biol. 18, 27–34.

Gertz, J., Reddy, T.E., Varley, K.E., Garabedian, M.J., and Myers, R.M. (2012). Genistein and bisphenol A exposure cause estrogen receptor 1 to bind thousands of sites in a cell type-specific manner. Genome research 22, 2153–2162.

Gertz, J., Savic, D., Varley, E., Katherine, Partridge C. E., Safi, A., Jain, P., Cooper, M., Gregory, Reddy, E., Timothy, Crawford E., Gregory, and Myers, M., Richard (2013). Distinct Properties of Cell-Type-Specific and Shared Transcription Factor Binding Sites. Mol. Cell 52, 25–36.

Ginley-Hidinger, M., Carleton, J.B., Rodriguez, A.C., Berrett, K.C., and Gertz, J. (2019). Sufficiency analysis of estrogen responsive enhancers using synthetic activators. Life Science Alliance 2, e201900497.

Hah, N., Benner, C., Chong, L.-W., Yu, R.T., Downes, M., and Evans, R.M. (2015). Inflammation-sensitive super enhancers form domains of coordinately regulated enhancer RNAs. Proceedings of the National Academy of Sciences 112, E297–E302.

Ham, L., Jackson, M., and Stumpf, M.P. (2021). Pathway dynamics can delineate the sources of transcriptional noise in gene expression. Elife 10.

Han, R., Huang, G., Wang, Y., Xu, Y., Hu, Y., Jiang, W., Wang, T., Xiao, T., and Zheng, D. (2016). Increased gene expression noise in human cancers is correlated with low p53 and immune activities as well as late stage cancer. Oncotarget 7, 72011–72020.

Hannon, G.J. (2010). FASTX-Toolkit FASTQ/A short-reads pre-processing tools, http://hannonlab.cshl.edu/fastx_toolkit/.

Hassig, C.A., Fleischer, T.C., Billin, A.N., Schreiber, S.L., and Ayer, D.E. (1997). Histone deacetylase activity is required for full transcriptional repression by mSin3A. Cell 89, 341–347.

Hnisz, D., Shrinivas, K., Young, R.A., Chakraborty, A.K., and Sharp, P.A. (2017). A Phase Separation Model for Transcriptional Control. Cell 169, 13–23.

Iyer, V.R., Eisen, M.B., Ross, D.T., Schuler, G., Moore, T., Lee, J.C., Trent, J.M., Staudt, L.M., Hudson, J., Jr., Boguski, M.S., et al. (1999). The transcriptional program in the response of human fibroblasts to serum. Science 283, 83–87.

Jagannathan, V., and Robinson-Rechavi, M. (2011). Meta-analysis of estrogen response in MCF-7 distinguishes early target genes involved in signaling and cell proliferation from later target genes involved in cell cycle and DNA repair. BMC systems biology 5, 1–14.

Juan, A.H., and Ruddle, F.H. (2003). Enhancer timing of Hox gene expression: deletion of the endogenous Hoxc8 early enhancer. Development 130, 4823–4834.

Kent, W.J., Sugnet, C.W., Furey, T.S., Roskin, K.M., Pringle, T.H., Zahler, A.M., and Haussler, D. (2002). The human genome browser at UCSC. Genome Res. 12, 996–1006.

Kim, D., Paggi, J.M., Park, C., Bennett, C., and Salzberg, S.L. (2019). Graph-based genome alignment and genotyping with HISAT2 and HISAT-genotype. Nat. Biotechnol. 37, 907–915.

Kolch, W., Halasz, M., Granovskaya, M., and Kholodenko, B.N. (2015). The dynamic control of signal transduction networks in cancer cells. Nature Reviews Cancer 15, 515–527.

Konstantinides, N., Holguera, I., Rossi, A.M., Escobar, A., Dudragne, L., Chen, Y.-C., Tran, T.N., Martínez Jaimes, A.M., Özel, M.N., Simon, F., et al. (2022). A complete temporal transcription factor series in the fly visual system. Nature 604, 316–322.

Krakauer, D.C., Page, K.M., and Sealfon, S. (2002). Module dynamics of the GnRH signal transduction network. J. Theor. Biol. 218, 457–470.

Kundaje, A., Meuleman, W., Ernst, J., Bilenky, M., Yen, A., Heravi-Moussavi, A., Kheradpour, P., Zhang, Z., Wang, J., Ziller, M.J., et al. (2015). Integrative analysis of 111 reference human epigenomes. Nature 518, 317–330.

Kursa, M.B., and Rudnicki, W.R. (2010a). Feature Selection with the Boruta Package. Journal of Statistical Software 36, 1–13.

Kursa, M.B., and Rudnicki, W.R. (2010b). Feature selection with the Boruta package. Journal of statistical software 36, 1–13.

Langmead, B., and Salzberg, S.L. (2012). Fast gapped-read alignment with Bowtie 2. Nat. Methods 9, 357–359.

Langmead, B., Trapnell, C., Pop, M., and Salzberg, S.L. (2009). Ultrafast and memory-efficient alignment of short DNA sequences to the human genome. Genome biology 10, 1–10.

Lareau, C.A., and Aryee, M.J. (2018). hichipper: a preprocessing pipeline for calling DNA loops from HiChIP data. Nat. Methods 15, 155–156.

Larsson, A.J.M., Johnsson, P., Hagemann-Jensen, M., Hartmanis, L., Faridani, O.R., Reinius, B., Segerstolpe, Å., Rivera, C.M., Ren, B., and Sandberg, R. (2019). Genomic encoding of transcriptional burst kinetics. Nature 565, 251–254.

Li, H., Handsaker, B., Wysoker, A., Fennell, T., Ruan, J., Homer, N., Marth, G., Abecasis, G., Durbin, R., and Subgroup, G.P.D.P. (2009). The Sequence Alignment/Map format and SAMtools. Bioinformatics 25, 2078–2079.

Liao, Y., Smyth, G.K., and Shi, W. (2014). featureCounts: an efficient general purpose program for assigning sequence reads to genomic features. Bioinformatics 30, 923–930.

Liberzon, A., Birger, C., Thorvaldsdóttir, H., Ghandi, M., Jill, and Tamayo, P. (2015). The Molecular Signatures Database Hallmark Gene Set Collection. Cell Systems 1, 417–425.

Love, M., Anders, S., and Huber, W. (2014). Differential analysis of count data–the DESeq2 package. Genome Biol 15, 10–1186.

MacKenzie, A., Hing, B., and Davidson, S. (2013). Exploring the effects of polymorphisms on cis-regulatory signal transduction response. Trends Mol. Med. 19, 99–107.

Mahat, D.B., Kwak, H., Booth, G.T., Jonkers, I.H., Danko, C.G., Patel, R.K., Waters, C.T., Munson, K., Core, L.J., and Lis, J.T. (2016). Base-pair-resolution genome-wide mapping of active RNA polymerases using precision nuclear run-on (PRO-seq). Nature Protocols 11, 1455–1476.

Martin, M. (2011). Cutadapt removes adapter sequences from high-throughput sequencing reads. 2011 *17*, 3.

McGinnis, C.S., Patterson, D.M., Winkler, J., Conrad, D.N., Hein, M.Y., Srivastava, V., Hu, J.L., Murrow, L.M., Weissman, J.S., Werb, Z., et al. (2019). MULTI-seq: sample multiplexing for single-cell RNA sequencing using lipid-tagged indices. Nat. Methods 16, 619–626.

Mumbach, M.R., Rubin, A.J., Flynn, R.A., Dai, C., Khavari, P.A., Greenleaf, W.J., and Chang, H.Y. (2016). HiChIP: efficient and sensitive analysis of protein-directed genome architecture. Nat. Methods 13, 919–922.

Murai, J., Zhang, H., Pongor, L., Tang, S.-W., Jo, U., Moribe, F., Ma, Y., Tomita, M., and Pommier, Y. (2020). Chromatin Remodeling and Immediate Early Gene Activation by SLFN11 in Response to Replication Stress. Cell Reports 30, 4137–4151.e4136.

Nguyen, A., Yoshida, M., Goodarzi, H., and Tavazoie, S.F. (2016). Highly variable cancer subpopulations that exhibit enhanced transcriptome variability and metastatic fitness. Nature communications 7, 1–13.

Nicolas, D., Zoller, B., Suter, D.M., and Naef, F. (2018). Modulation of transcriptional burst frequency by histone acetylation. Proceedings of the National Academy of Sciences 115, 7153–7158.

Ostuni, R., Piccolo, V., Barozzi, I., Polletti, S., Termanini, A., Bonifacio, S., Curina, A., Prosperini, E., Ghisletti, S., and Natoli, G. (2013). Latent Enhancers Activated by Stimulation in Differentiated Cells. Cell 152, 157–171.

Parab, L., Pal, S., and Dhar, R. (2022). Transcription factor binding process is the primary driver of noise in gene expression. PLoS Genet. 18, e1010535.

Pedraza, J.M., Garcia, D.A., and Pérez-Ortiz, M.F. (2018). Noise, information and fitness in changing environments. Frontiers in Physics 6, 83.

Pruitt, K.D., Brown, G.R., Hiatt, S.M., Thibaud-Nissen, F., Astashyn, A., Ermolaeva, O., Farrell, C.M., Hart, J., Landrum, M.J., McGarvey, K.M., et al. (2013). RefSeq: an update on mammalian reference sequences. Nucleic Acids Res. 42, D756–D763.

Qin, S., Jiang, J., Lu, Y., Nice, E.C., Huang, C., Zhang, J., and He, W. (2020). Emerging role of tumor cell plasticity in modifying therapeutic response. Signal Transduct Target Ther 5, 228.

Raj, A., and Van Oudenaarden, A. (2008). Nature, Nurture, or Chance: Stochastic Gene Expression and Its Consequences. Cell 135, 216–226.

Raser, J.M., and O’Shea, E.K. (2005). Noise in Gene Expression: Origins, Consequences, and Control. Science 309, 2010–2013.

Reddy, T.E., Pauli, F., Sprouse, R.O., Neff, N.F., Newberry, K.M., Garabedian, M.J., and Myers, R.M. (2009). Genomic determination of the glucocorticoid response reveals unexpected mechanisms of gene regulation. Genome Res. 19, 2163–2171.

Rodriguez, A.C., Blanchard, Z., Maurer, K.A., and Gertz, J. (2019a). Estrogen Signaling in Endometrial Cancer: a Key Oncogenic Pathway with Several Open Questions. Hormones and Cancer 10, 51–63.

Rodriguez, J., Ren, G., Day, C.R., Zhao, K., Chow, C.C., and Larson, D.R. (2019b). Intrinsic Dynamics of a Human Gene Reveal the Basis of Expression Heterogeneity. Cell 176, 213–226 e218.

Satija, R., Farrell, J.A., Gennert, D., Schier, A.F., and Regev, A. (2015). Spatial reconstruction of single-cell gene expression data. Nat. Biotechnol. 33, 495–502.

Schier, A.C., and Taatjes, D.J. (2020). Structure and mechanism of the RNA polymerase II transcription machinery. Genes & Development 34, 465–488.

Schnoes, K.K., Jaffe, I.Z., Iyer, L., Dabreo, A., Aronovitz, M., Newfell, B., Hansen, U., Rosano, G., and Mendelsohn, M.E. (2008). Research Resource: Rapid Recruitment of Temporally Distinct Vascular Gene Sets by Estrogen. Mol. Endocrinol. 22, 2544–2556.

Servant, N., Varoquaux, N., Lajoie, B.R., Viara, E., Chen, C.-J., Vert, J.-P., Heard, E., Dekker, J., and Barillot, E. (2015). HiC-Pro: an optimized and flexible pipeline for Hi-C data processing. Genome biology 16, 1–11.

Shaffer, S.M., Dunagin, M.C., Torborg, S.R., Torre, E.A., Emert, B., Krepler, C., Beqiri, M., Sproesser, K., Brafford, P.A., Xiao, M., et al. (2017). Rare cell variability and drug-induced reprogramming as a mode of cancer drug resistance. Nature 546, 431–435.

Sheng, M., and Greenberg, M.E. (1990). The regulation and function of c-fos and other immediate early genes in the nervous system. Neuron 4, 477–485.

Shu, S., Lin, C.Y., He, H.H., Witwicki, R.M., Tabassum, D.P., Roberts, J.M., Janiszewska, M., Jin Huh, S., Liang, Y., Ryan, J., et al. (2016). Response and resistance to BET bromodomain inhibitors in triple-negative breast cancer. Nature 529, 413–417.

Shvartsur, A., and Bonavida, B. (2014). Trop2 and its overexpression in cancers: regulation and clinical/ therapeutic implications. Genes & Cancer 6, 84–105.

Sigalova, O.M., Shaeiri, A., Forneris, M., Furlong, E.E., and Zaugg, J.B. (2020). Predictive features of gene expression variation reveal mechanistic link with differential expression. Mol. Syst. Biol. 16, e9539.

Simeonov, D.R., Gowen, B.G., Boontanrart, M., Roth, T.L., Gagnon, J.D., Mumbach, M.R., Satpathy, A.T., Lee, Y., Bray, N.L., Chan, A.Y., et al. (2017). Discovery of stimulation-responsive immune enhancers with CRISPR activation. Nature 549, 111–115.

Stanford, J.L., Szklo, M., and Brinton, L.A. (1986). Estrogen receptors and breast cancer. Epidemiol. Rev. 8, 42–59.

Stuart, T., Butler, A., Hoffman, P., Hafemeister, C., Papalexi, E., Mauck, W.M., Hao, Y., Stoeckius, M., Smibert, P., and Satija, R. (2019). Comprehensive Integration of Single-Cell Data. Cell 177, 1888–1902.e1821.

Suderman, R., Bachman, J.A., Smith, A., Sorger, P.K., and Deeds, E.J. (2017). Fundamental trade-offs between information flow in single cells and cellular populations. Proceedings of the National Academy of Sciences 114, 5755–5760.

Szustakowski, J.D., Kosinski, P.A., Marrese, C.A., Lee, J.-H., Elliman, S.J., Nirmala, N., and Kemp, D.M. (2007). Dynamic resolution of functionally related gene sets in response to acute heat stress. BMC Mol. Biol. 8, 46.

Tani, H., Mizutani, R., Salam, K.A., Tano, K., Ijiri, K., Wakamatsu, A., Isogai, T., Suzuki, Y., and Akimitsu, N. (2012). Genome-wide determination of RNA stability reveals hundreds of short-lived noncoding transcripts in mammals. Genome Res. 22, 947–956.

Team, R.C. (2013). R: A language and environment for statistical computing.

Tullai, J.W., Schaffer, M.E., Mullenbrock, S., Sholder, G., Kasif, S., and Cooper, G.M. (2007). Immediate-Early and Delayed Primary Response Genes Are Distinct in Function and Genomic Architecture. J. Biol. Chem. 282, 23981–23995.

Uhlitz, F., Sieber, A., Wyler, E., FritscheLGuenther, R., Meisig, J., Landthaler, M., Klinger, B., and Blüthgen, N. (2017). An immediate–late gene expression module decodes ERK signal duration. Mol. Syst. Biol. 13, 928.

Urban, E.A., and Johnston, R.J., Jr. (2018). Buffering and Amplifying Transcriptional Noise During Cell Fate Specification. Front Genet 9, 591.

Uribe, M.L., Marrocco, I., and Yarden, Y. (2021). EGFR in cancer: Signaling mechanisms, drugs, and acquired resistance. Cancers 13, 2748.

Urrutia, R. (2003). KRAB-containing zinc-finger repressor proteins. Genome Biol 4, 231.

Wang, Z., Zang, C., Rosenfeld, J.A., Schones, D.E., Barski, A., Cuddapah, S., Cui, K., Roh, T.-Y., Peng, W., Zhang, M.Q., et al. (2008). Combinatorial patterns of histone acetylations and methylations in the human genome. Nat. Genet. 40, 897–903.

Wei, Y., Gokhale, R.H., Sonnenschein, A., Montgomery, K.M.t., Ingersoll, A., and Arnosti, D.N. (2016). Complex cis-regulatory landscape of the insulin receptor gene underlies the broad expression of a central signaling regulator. Development 143, 3591–3603.

Wibisana, J.N., Inaba, T., Shinohara, H., Yumoto, N., Hayashi, T., Umeda, M., Ebisawa, M., Nikaido, I., Sako, Y., and Okada, M. (2022). Enhanced transcriptional heterogeneity mediated by NF-κB super-enhancers. PLoS Genet. 18, e1010235.

Wollman, R. (2018). Robustness, Accuracy, and Cell State Heterogeneity in Biological Systems. Curr Opin Syst Biol 8, 46–50.

Wood, S.N. (2006). Generalized additive models: an introduction with R (chapman and hall/CRC).

Wu, S., Li, K., Li, Y., Zhao, T., Li, T., Yang, Y.-F., and Qian, W. (2017). Independent regulation of gene expression level and noise by histone modifications. PLoS Comp. Biol. 13, e1005585.

Xie, Z., Bailey, A., Kuleshov, M.V., Clarke, D.J., Evangelista, J.E., Jenkins, S.L., Lachmann, A., Wojciechowicz, M.L., Kropiwnicki, E., and Jagodnik, K.M. (2021). Gene set knowledge discovery with Enrichr. Current protocols 1, e90.

Zhang, G., Zhao, Y., Liu, Y., Kao, L.-P., Wang, X., Skerry, B., and Li, Z. (2016). FOXA1 defines cancer cell specificity. Science Advances 2, e1501473.

Zhang, J., Lee, D., Dhiman, V., Jiang, P., Xu, J., McGillivray, P., Yang, H., Liu, J., Meyerson, W., Clarke, D., et al. (2020). An integrative ENCODE resource for cancer genomics. Nature Communications 11.

Zhang, J., Zhu, W., Wang, Q., Gu, J., Huang, L.F., and Sun, X. (2019). Differential regulatory network-based quantification and prioritization of key genes underlying cancer drug resistance based on time-course RNA-seq data. PLoS Comp. Biol. 15, e1007435.

Zhang, Y., Liu, T., Meyer, C.A., Eeckhoute, J., Johnson, D.S., Bernstein, B.E., Nusbaum, C., Myers, R.M., Brown, M., and Li, W. (2008). Model-based analysis of ChIP-Seq (MACS). Genome biology 9, 1–9.

