## Supplementary figures and tables for "*Cis*-regulatory control of transcriptional timing and noise in response to estrogen"

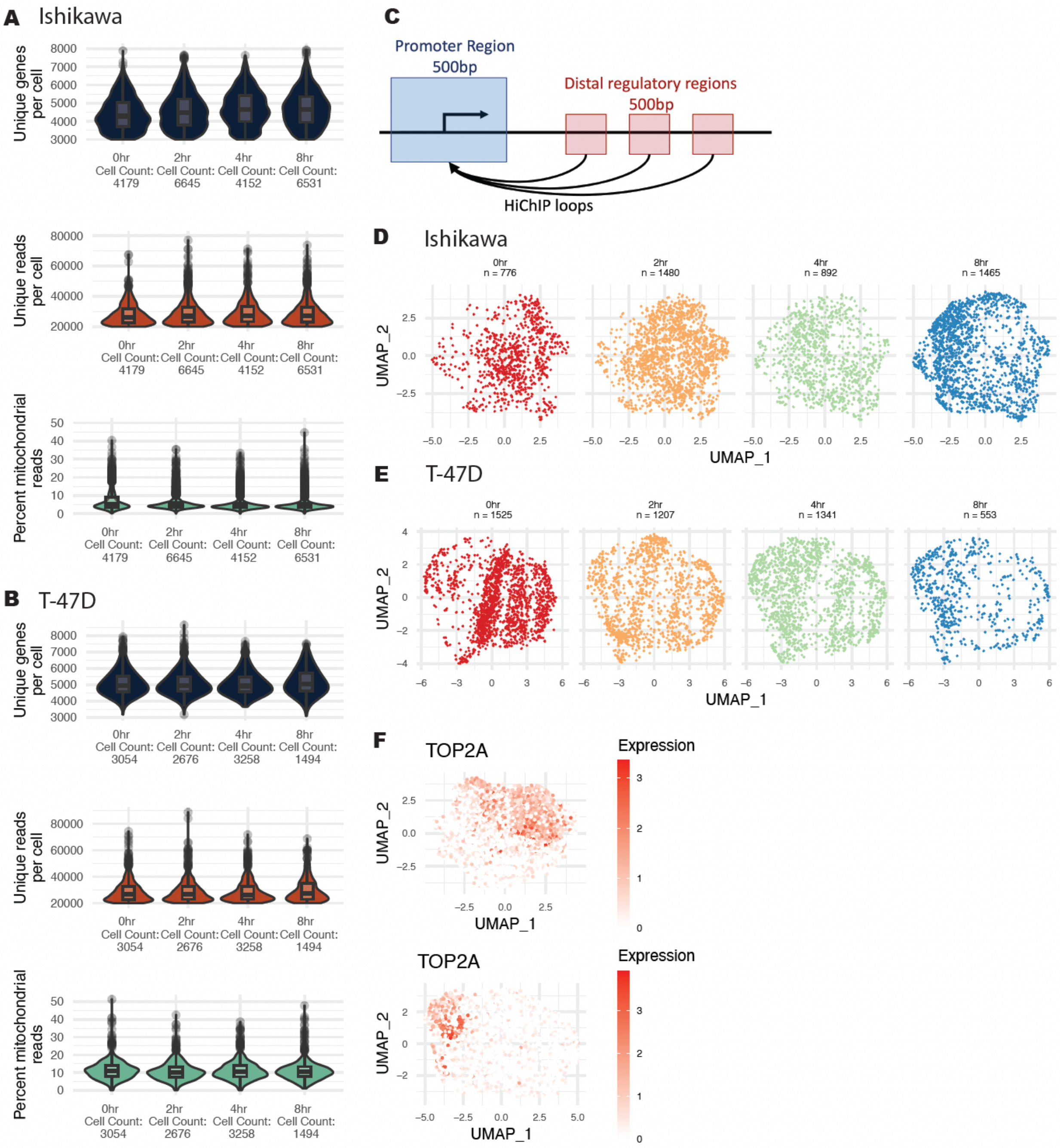

**Supplementary Figure 1 (related to Figures 1 and 2). QC plots for scRNA-seq and diagram of *cis*-regulatory model.** (A-B) Distribution of reads, genes, and percent mitochondrial reads per cell, separated by timepoint, are shown for (A) Ishikawa and (B) T-47D. Data shown is prior to filtering. (C) A model shows how signal intensity was collected for promoters and enhancers using HiChIP loops to identify enhancers. (D-E) UMAP dimensionality reduction plots for (D) Ishikawa and (E) T-47D show temporal progression of cells treated with E2. Each point represents a cell and timepoints are post 10 nM E2 induction. Cell numbers represent post-filtering. All cells for a cell line were used to create the original UMAP projections and the space was held constant when plotting each time point. (F) UMAP representation of cell-cycle marker gene *TOP2A,* which peaks in the G2/M phase, is shown to highlight the location of proliferating cells.

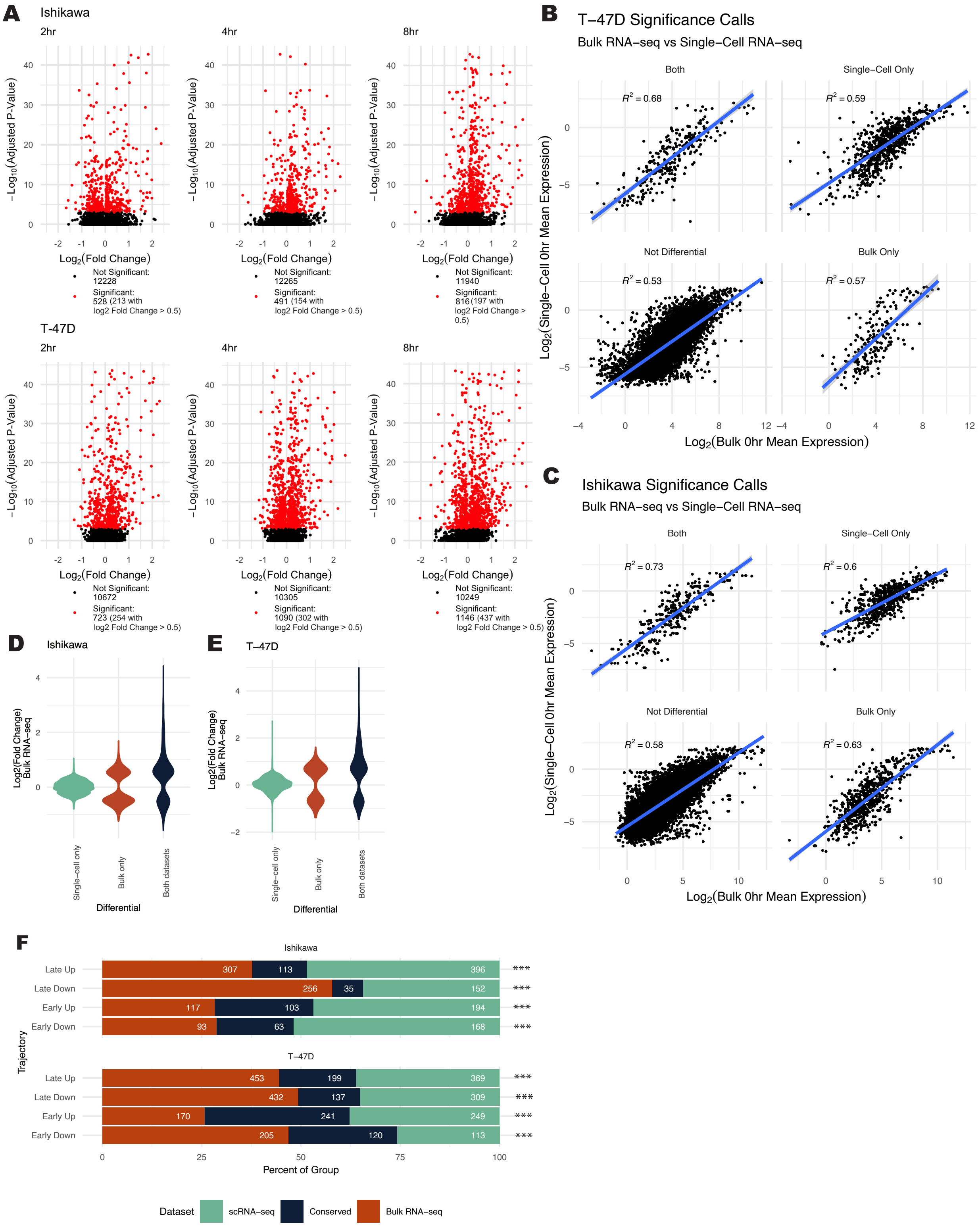
A
Ishikawa
2hr
4hr
8hr
−Log10(Adjusted P-Value)
Log2(Fold Change)
Not Significant: 12228
Significant: 528 (213 with log2 Fold Change > 0.5)
Not Significant: 12265
Significant: 491 (154 with log2 Fold Change > 0.5)
Not Significant: 11940
Significant: 816 (197 with log2 Fold Change > 0.5)
T-47D
Not Significant: 10672
Significant: 723 (254 with log2 Fold Change > 0.5)
Not Significant: 10305
Significant: 1090 (302 with log2 Fold Change > 0.5)
Not Significant: 10249
Significant: 1146 (437 with log2 Fold Change > 0.5)
B
T−47D Significance Calls
Bulk RNA−seq vs Single−Cell RNA−seq
Both
Single−Cell Only
Not Differential
Bulk Only
R2 = 0.68
R2 = 0.59
R2 = 0.53
R2 = 0.57
Log2(Single−Cell 0hr Mean Expression)
Log2(Bulk 0hr Mean Expression)
C
Ishikawa Significance Calls
Bulk RNA−seq vs Single−Cell RNA−seq
Both
Single−Cell Only
Not Differential
Bulk Only
R2 = 0.73
R2 = 0.6
R2 = 0.58
R2 = 0.63
Log2(Single−Cell 0hr Mean Expression)
Log2(Bulk 0hr Mean Expression)
D
Ishikawa
E
T−47D
Log2(Fold Change) Bulk RNA−seq
Single−cell only
Bulk only
Both datasets
Differential
F
Ishikawa
T−47D
Late Up
Late Down
Early Up
Early Down
Trajectory
307 113 396 ***
256 35 152 ***
117 103 194 ***
93 63 168 ***
453 199 369 ***
432 137 309 ***
170 241 249 ***
205 120 113 ***
Percent of Group
Dataset
scRNA−seq
Conserved
Bulk RNA−seq

**Supplementary Figure 2 (related to Figure 2). Analysis of differential gene expression over time.** (A) Differential gene expression -$\log_{10}$(Wilcoxon p-values) are plotted against $\log_2$(fold change) for all genes at each timepoint vs 0 hours. Significantly differential genes are highlighted in red. Data is shown for Ishikawa (top) and T-47D (bottom) cells. (B-C) Average Reads Per Million from scRNA-seq are correlated to bulk RNA-seq RPM in both Ishikawa (B) and T-47D (C) cells, shown on a log scale. Genes are separated into groups depending on whether they are called significant in bulk RNA-seq, scRNA-seq, or both. (D-E) $\mathrm{Log}_2$ fold change distribution is shown for differential genes called by scRNA-seq analysis or bulk RNA-seq analysis in Ishikawa (D) and T-47D (E) cells. (F) Trajectory analysis in bulk RNA-seq vs scRNA-seq reveals a significant overlap of genes classified into each trajectory. P-values are calculated using hypergeometric tests (***: $p < 1 \times 10^{-28}$).

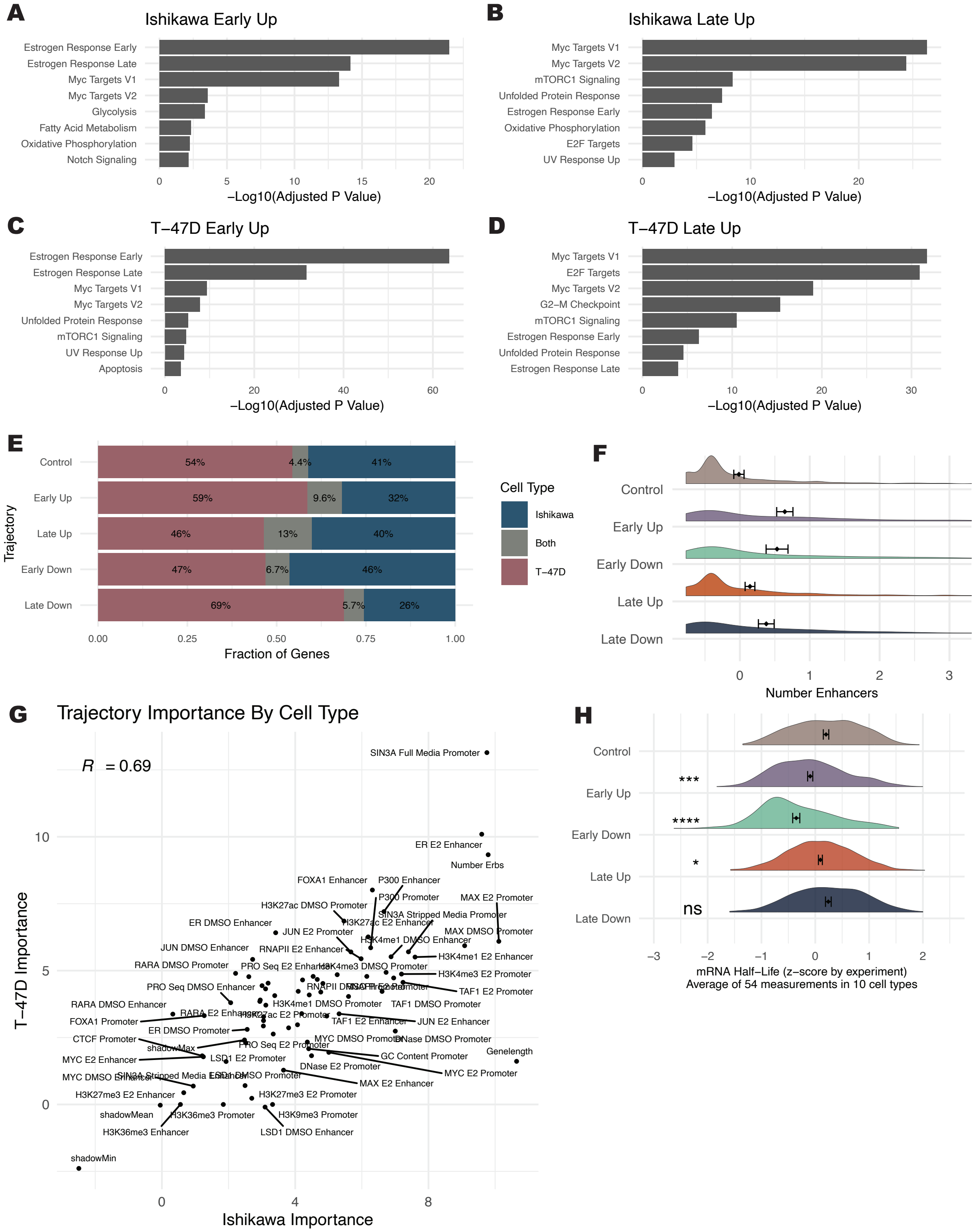

A
Ishikawa Early Up
Estrogen Response Early
Estrogen Response Late
Myc Targets V1
Myc Targets V2
Glycolysis
Fatty Acid Metabolism
Oxidative Phosphorylation
Notch Signaling
0 5 10 15 20
−Log10(Adjusted P Value)
B
Ishikawa Late Up
Myc Targets V1
Myc Targets V2
mTORC1 Signaling
Unfolded Protein Response
Estrogen Response Early
Oxidative Phosphorylation
E2F Targets
UV Response Up
0 10 20
−Log10(Adjusted P Value)
C
T−47D Early Up
Estrogen Response Early
Estrogen Response Late
Myc Targets V1
Myc Targets V2
Unfolded Protein Response
mTORC1 Signaling
UV Response Up
Apoptosis
0 20 40 60
−Log10(Adjusted P Value)
D
T−47D Late Up
Myc Targets V1
E2F Targets
Myc Targets V2
G2−M Checkpoint
mTORC1 Signaling
Estrogen Response Early
Unfolded Protein Response
Estrogen Response Late
0 10 20 30
−Log10(Adjusted P Value)
E
Trajectory
Control 54% 4.4% 41%
Early Up 59% 9.6% 32%
Late Up 46% 13% 40%
Early Down 47% 6.7% 46%
Late Down 69% 5.7% 26%
0.00 0.25 0.50 0.75 1.00
Fraction of Genes
Cell Type
Ishikawa
Both
T−47D
F
Control
Early Up
Early Down
Late Up
Late Down
0 1 2 3
Number Enhancers
G
Trajectory Importance By Cell Type
R = 0.69
SIN3A Full Media Promoter
ER E2 Enhancer
Number Erbs
FOXA1 Enhancer
P300 Enhancer
P300 Promoter
MAX E2 Promoter
H3K27ac DMSO Promoter
H3K27ac E2 Enhancer
SIN3A Stripped Media Promoter
ER DMSO Enhancer
JUN E2 Promoter
MAX DMSO Promoter
H3K4me1 DMSO Enhancer
JUN DMSO Enhancer
RNAPII E2 Enhancer
H3K4me1 E2 Enhancer
RARA DMSO Promoter
PRO Seq E2 Enhancer
H3K4me3 DMSO Promoter
H3K4me3 E2 Promoter
PRO Seq DMSO Enhancer
TAF1 E2 Promoter
RARA DMSO Enhancer
H3K4me1 DMSO Promoter
TAF1 DMSO Promoter
FOXA1 Promoter
RARA E2 Enhancer
H3K27ac E2 Promoter
TAF1 E2 Enhancer
JUN E2 Enhancer
ER DMSO Promoter
CTCF Promoter
MYC DMSO Promoter
DNase DMSO Promoter
shadowMax
PRO Seq E2 Promoter
Genelength
MYC E2 Enhancer
LSD1 E2 Promoter
GC Content Promoter
DNase E2 Promoter
MYC E2 Promoter
MYC DMSO Enhancer
SIN3A Stripped Media Enhancer
LSD1 DMSO Promoter
MAX E2 Enhancer
H3K27me3 E2 Enhancer
H3K27me3 E2 Promoter
shadowMean
H3K36me3 Promoter
H3K9me3 Promoter
H3K36me3 Enhancer
LSD1 DMSO Enhancer
shadowMin
T−47D Importance
0 5 10
Ishikawa Importance
0 4 8
H
Control
Early Up ***
Early Down ****
Late Up *
Late Down ns
−3 −2 −1 0 1 2
mRNA Half−Life (z−score by experiment)
Average of 54 measurements in 10 cell types

**Supplementary Figure 3 (related to Figure 2).** (A-D) Gene set enrichment of Early Up and Late Up genes for MSigDB Hallmark 2020 is shown for Ishikawa (A-B) and T-47D (C-D) cells. P-values are calculated using EnrichR. (E) Proportion of overlap between genes of each trajectory classification between cell types is shown. (F) Distribution in number of enhancers (z-score) by trajectory is plotted. Data is combined from both Ishikawa and T-47D cells. (G) The scatterplot shows median Boruta importance values for classifying trajectory in T-47D (y-axis) vs Ishikawa (x-axis). Boruta analysis was run separately in each cell type. Each data point represents a genomic feature. The correlation value was calculated using Pearson correlation. (H) mRNA half-life values from across 54 human samples were converted to Z-scores within each experiment. An average value was plotted for each gene, classified into trajectories based on the timing of the transcriptional response to E2. p-values are calculated using a Kolmogorov-Smirnov Test vs the Control group (ns > 0.05, * < 0.05, ** < 1e-5, *** < 1e-10, **** < 1e-15).

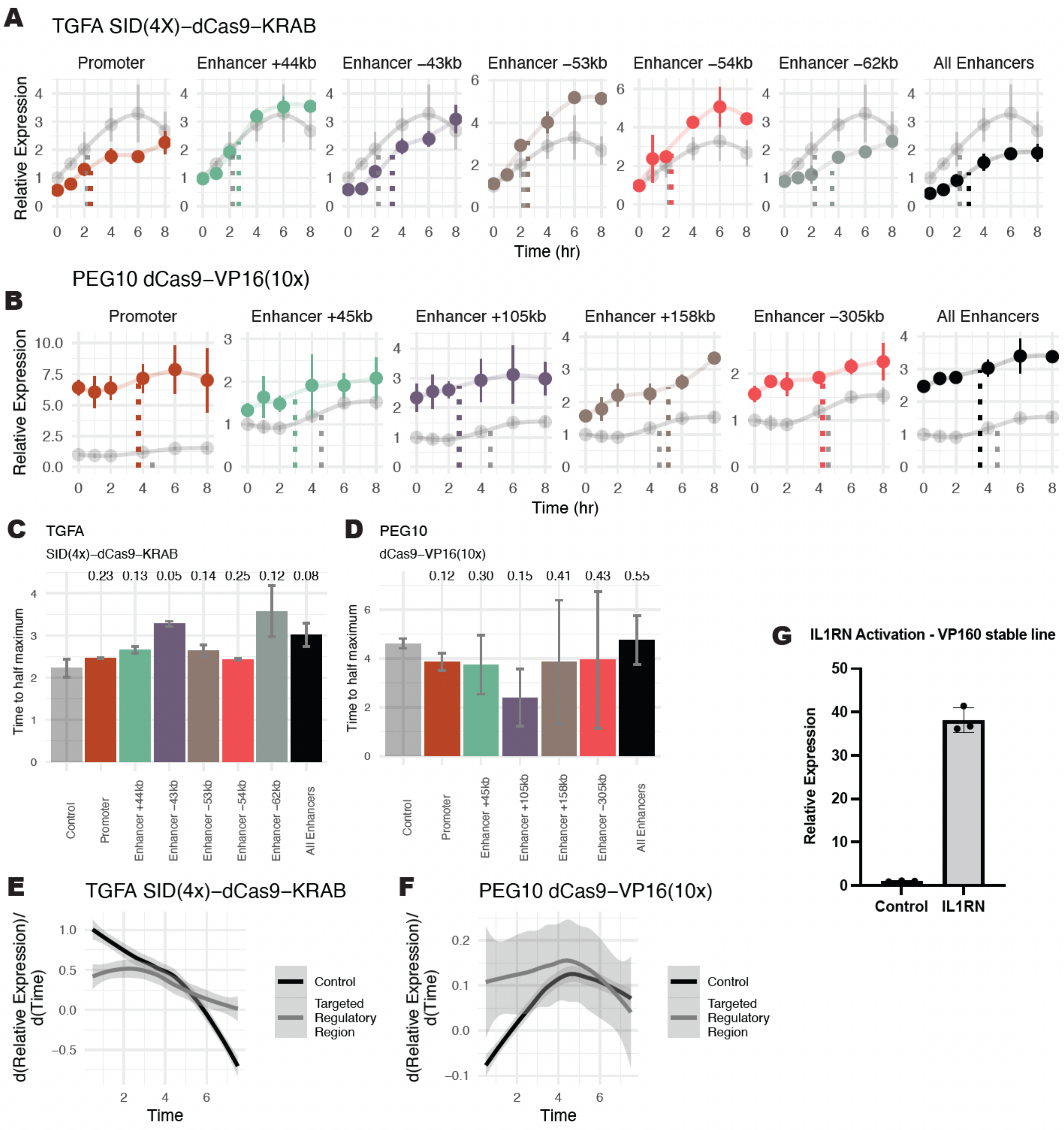
A
TGFA SID(4X)–dCas9–KRAB
Promoter
Enhancer +44kb
Enhancer −43kb
Enhancer −53kb
Enhancer −54kb
Enhancer −62kb
All Enhancers
Relative Expression
Time (hr)
B
PEG10 dCas9–VP16(10x)
Promoter
Enhancer +45kb
Enhancer +105kb
Enhancer +158kb
Enhancer −305kb
All Enhancers
Relative Expression
Time (hr)
C
TGFA
SID(4x)–dCas9–KRAB
0.23 0.13 0.05 0.14 0.25 0.12 0.08
Time to half maximum
Control
Promoter
Enhancer +44kb
Enhancer −43kb
Enhancer −53kb
Enhancer −54kb
Enhancer −62kb
All Enhancers
D
PEG10
dCas9–VP16(10x)
0.12 0.30 0.15 0.41 0.43 0.55
Time to half maximum
Control
Promoter
Enhancer +45kb
Enhancer +105kb
Enhancer +158kb
Enhancer −305kb
All Enhancers
G
IL1RN Activation - VP160 stable line
Relative Expression
Control
IL1RN
E
TGFA SID(4x)–dCas9–KRAB
d(Relative Expression)/d(Time)
Time
Control
Targeted Regulatory Region
F
PEG10 dCas9–VP16(10x)
d(Relative Expression)/d(Time)
Time
Control
Targeted Regulatory Region

**Supplementary Figure 4 (related to Figure 3). Synthetic transcription factors targeted to regulatory regions alter trajectories of *TGFA* and *PEG10*.** (A) Expression trajectory of *TGFA* following E2 induction in Ishikawa cells is shown. Regulatory regions are targeted with SID(4x)-dCas9-KRAB. Data from cells with *IL1RN* promoter targeting (control) are shown in gray. Dotted lines represent the time to half maximal expression for each trajectory. Expression is relative to the 0-hour timepoint in the control. (B) *PEG10* gene expression trajectories in Ishikawa cells following E2 inductions are modulated by dCas9-VP16(10x) targeted to regulatory regions. Dotted lines represent the time to half maximal expression for each trajectory. Expression is relative to the 0-hour timepoint in the control. (C-D) Bar plot shows time to half maximal expression for each targeted regulatory region. Error bars represent the SEM (n=2) and p-values (one-sided t-test) are reported above each bar. (E-F) Aggregate differential of loess regressions from panels A and B show expression slope when CREs near *TGFA* are targeted by SID(4x)-dCas9-KRAB (E) or *PEG10* CREs are targeted by dCas9-VP16(10x) (F). Slopes are aggregated targeted (grey) compared to control (black) and the shaded region represents 95% confidence interval. (G) Validation of stable dCas9-VP16(10x) expressing Ishikawa cell line shows a control gene, *IL1RN,* being activated to previously observed levels (Ginley-Hidinger et al., 2019).

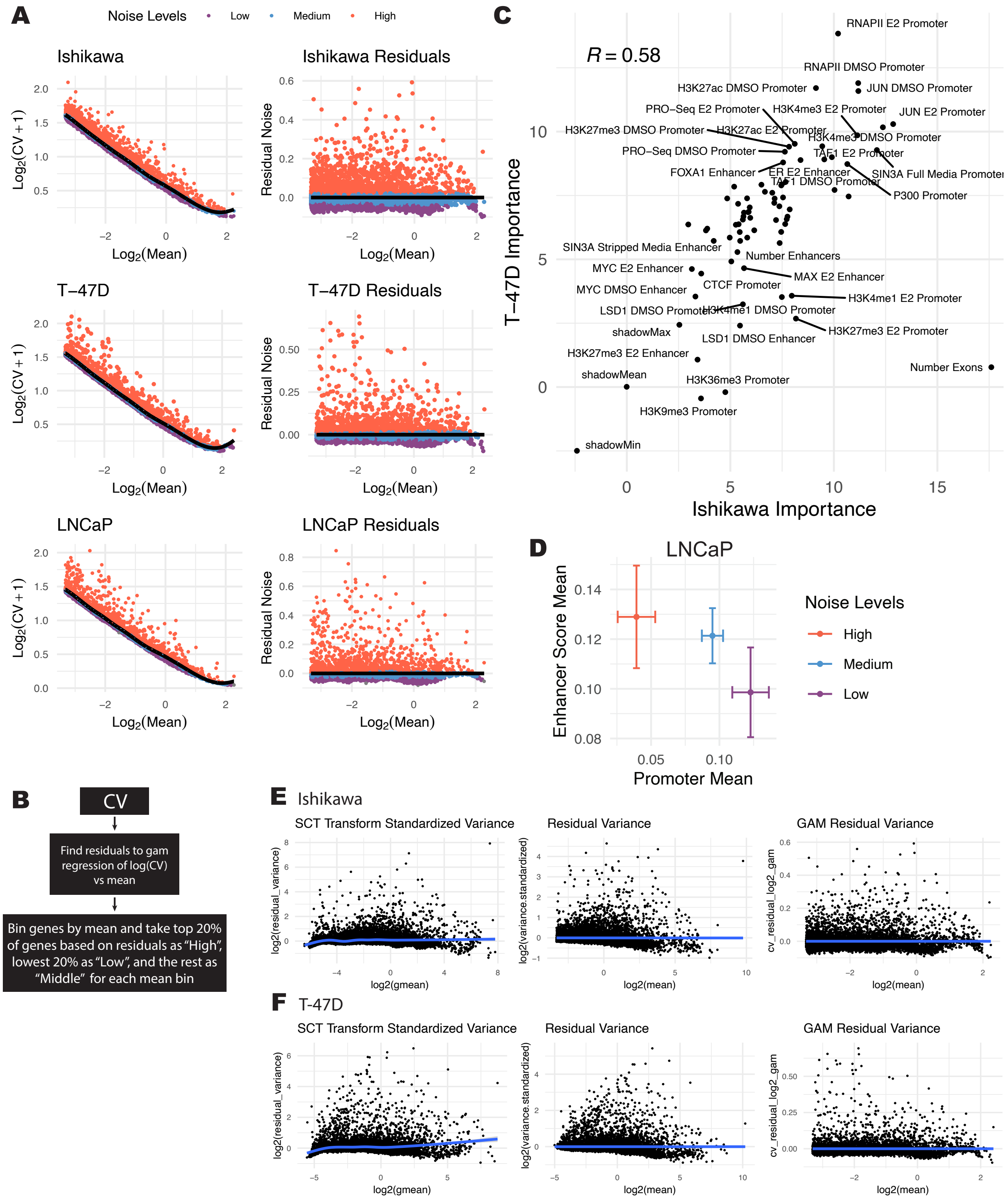
A
Noise Levels
Low
Medium
High
Ishikawa
Ishikawa Residuals
T−47D
T−47D Residuals
LNCaP
LNCaP Residuals
Log2(CV + 1)
Residual Noise
Log2(Mean)
B
CV
Find residuals to gam regression of log(CV) vs mean
Bin genes by mean and take top 20% of genes based on residuals as "High", lowest 20% as "Low", and the rest as "Middle" for each mean bin
C
R = 0.58
T−47D Importance
Ishikawa Importance
RNAPII E2 Promoter
RNAPII DMSO Promoter
H3K27ac DMSO Promoter
JUN DMSO Promoter
PRO−Seq E2 Promoter
H3K4me3 E2 Promoter
JUN E2 Promoter
H3K27me3 DMSO Promoter
H3K27ac E2 Promoter
H3K4me3 DMSO Promoter
PRO−Seq DMSO Promoter
TAF1 E2 Promoter
FOXA1 Enhancer
ER E2 Enhancer
SIN3A Full Media Promoter
TAF1 DMSO Promoter
P300 Promoter
SIN3A Stripped Media Enhancer
Number Enhancers
MYC E2 Enhancer
MAX E2 Enhancer
CTCF Promoter
MYC DMSO Enhancer
H3K4me1 E2 Promoter
LSD1 DMSO Promoter
H3K4me1 DMSO Promoter
shadowMax
H3K27me3 E2 Promoter
LSD1 DMSO Enhancer
H3K27me3 E2 Enhancer
Number Exons
shadowMean
H3K36me3 Promoter
H3K9me3 Promoter
shadowMin
D
LNCaP
Enhancer Score Mean
Promoter Mean
Noise Levels
High
Medium
Low
E
Ishikawa
SCT Transform Standardized Variance
Residual Variance
GAM Residual Variance
log2(residual_variance)
log2(variance.standardized)
cv_residual_log2_gam
log2(gmean)
log2(mean)
F
T-47D
SCT Transform Standardized Variance
Residual Variance
GAM Residual Variance

**Supplementary Figure 5 (related to Figure 4). Strong enhancers and weak promoters are associated with high levels of noise.** (A) Calculated coefficient of variation (CV) is shown before (left panel) and after (right panel) adjustment using a generalized additive model regression fit for each cell type. Axes are on a log scale. Colors represent classified noise levels. (B) A flowchart depicts the regularization strategy to remove mean effects from the measured variation and to classify genes into noise levels. (C) Boruta importance scores for classifying genes into noise categories are shown for independent analyses in T-47D (y-axis) and Ishikawa (x-axis) cells. Correlation value was calculated using Pearson correlation. (D) Enhancer score averages vs promoter score averages are shown for noise levels in LNCaP cells. Averages are based on confirmed variables from a Boruta analysis. Axes do not represent full range. For a list of genomic data used in LNCaP cells see Table S3. (E-F) A comparison of noise regularization strategies in (E) Ishikawa and (F) T-47D cells is shown. SCT transform standardized variance and residual variance were calculated using the Seurat package in R. Blue line shows a loess best fit.

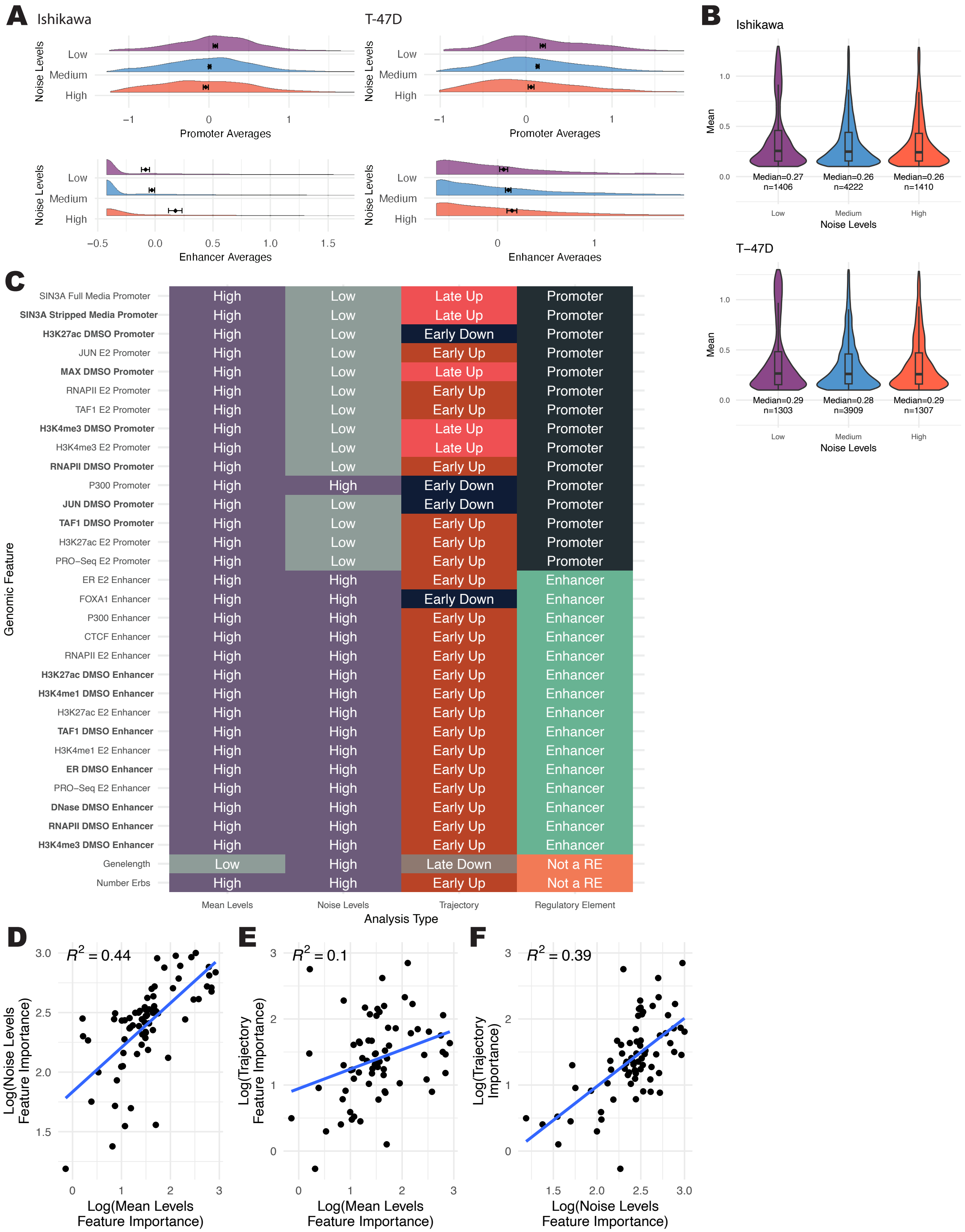
A
Ishikawa
T-47D
Noise Levels
Low
Medium
High
Promoter Averages
Enhancer Averages
B
Ishikawa
Mean
Median=0.27
n=1406
Median=0.26
n=4222
Median=0.26
n=1410
Noise Levels
T−47D
Median=0.29
n=1303
Median=0.28
n=3909
Median=0.29
n=1307
C
Genomic Feature
SIN3A Full Media Promoter
SIN3A Stripped Media Promoter
H3K27ac DMSO Promoter
JUN E2 Promoter
MAX DMSO Promoter
RNAPII E2 Promoter
TAF1 E2 Promoter
H3K4me3 DMSO Promoter
H3K4me3 E2 Promoter
RNAPII DMSO Promoter
P300 Promoter
JUN DMSO Promoter
TAF1 DMSO Promoter
H3K27ac E2 Promoter
PRO−Seq E2 Promoter
ER E2 Enhancer
FOXA1 Enhancer
P300 Enhancer
CTCF Enhancer
RNAPII E2 Enhancer
H3K27ac DMSO Enhancer
H3K4me1 DMSO Enhancer
H3K27ac E2 Enhancer
TAF1 DMSO Enhancer
H3K4me1 E2 Enhancer
ER DMSO Enhancer
PRO−Seq E2 Enhancer
DNase DMSO Enhancer
RNAPII DMSO Enhancer
H3K4me3 DMSO Enhancer
Genelength
Number Erbs
Mean Levels
Noise Levels
Trajectory
Regulatory Element
Analysis Type
High
Low
Late Up
Early Up
Early Down
Late Down
Promoter
Enhancer
Not a RE
D
$R^2 = 0.44$
Log(Noise Levels Feature Importance)
Log(Mean Levels Feature Importance)
E
$R^2 = 0.1$
Log(Trajectory Feature Importance)
Log(Mean Levels Feature Importance)
F
$R^2 = 0.39$
Log(Trajectory Importance)
Log(Noise Levels Feature Importance)

**Supplementary Figure 6 (related to Figure 5). Promoters and enhancers underlie different relationships between mean, noise, and trajectory.** (A) The full distribution of promoter (top) and enhancer (bottom) in Ishikawa (left) and T-47D (right) across noise levels is shown. (B) Graph depicts mean levels across different noise levels for Ishikawa (top) and T-47D (bottom). (C) Top 15 most important features for each regulatory element type (promoter or enhancer) are shown, determined by average rank across all three analysis types. For each analysis, the classification group with the highest signal is displayed. Datasets shown in bold were performed in the absence of ER activation. (D-F) Pairwise correlation plots show the relationship between importance scores for the three analysis types. Each point represents a genomic feature. $R^2$ values were calculated using Spearman correlation.

**Table S1. gRNA sequences**

| gRNA Name | Sequence | PAM |
|---|---|---|
| PEG10 Promoter gRNA 1 | GCCTTATATAGGTTGCACCG | AGG |
| PEG10 Promoter gRNA 2 | CGCGTGGTGAGTATGCGGTG | AGG |
| PEG10 Promoter gRNA 3 | GGAGTACGGGATTACCGAGG | GGG |
| PEG10 Promoter gRNA 4 | TTAGTGGCCTGCGCGCGACG | TGG |
| PEG10 Enhancer +105kb gRNA 1 | CATCTCCAGTTCAAACAGTG | AGG |
| PEG10 Enhancer +105kb gRNA 2 | AAGCCCAGGTATATTAATGG | AGG |
| PEG10 Enhancer +105kb gRNA 3 | TGACATAAACTAGCTGAGCA | GGG |
| PEG10 Enhancer +105kb gRNA 4 | ACTTGAGCTTGGATCCAGCT | GGG |
| PEG10 Enhancer +158kb gRNA 1 | GAGAGTTCCACTCTCGTGAA | TGG |
| PEG10 Enhancer +158kb gRNA 2 | GACAGAGCAAGACTCCGTAA | AGG |
| PEG10 Enhancer +158kb gRNA 3 | ATCTGCTGGAACCTTGATCA | TGG |
| PEG10 Enhancer +158kb gRNA 4 | GAGGAGAGCCTCTTGAACCT | GGG |
| PEG10 Enhancer +45kb gRNA 1 | GGAATATGCTCATAACCACA | AGG |
| PEG10 Enhancer +45kb gRNA 2 | CTCTTAATCTCCCTTCACCA | AGG |
| PEG10 Enhancer +45kb gRNA 3 | TCCTATTTCCTGTTGCCCTG | GGG |
| PEG10 Enhancer +45kb gRNA 4 | CCTAATTCCAGGCTCACCAC | AGG |
| PEG10 Enhancer +-305kb gRNA 1 | GGCATACTGGAATAAACATG | AGG |
| PEG10 Enhancer +-305kb gRNA 2 | TGAGAGACTCACTTGTCACT | AGG |
| PEG10 Enhancer +-305kb gRNA 3 | ACACTTCTAGAAGGCCCACT | AGG |
| PEG10 Enhancer +-305kb gRNA 4 | TAAAGAAGGTATAAGATCCA | TGG |
| TACSTD2 Promoter gRNA 1 | GGTAGAGTATAAGAGCCGGA | GGG |
| TACSTD2 Promoter gRNA 2 | TGTTCTGATCCTATCGCGGG | CGG |
| TACSTD2 Promoter gRNA 3 | GAGAACCGATACTCACCTGT | AGG |
| TACSTD2 Promoter gRNA 4 | TAACACCAGTCCTGACAGGT | AGG |
| TACSTD2 Enhancer +4.7kb gRNA 1 | CATCAGGAGACTAACCAGGG | AGG |
| TACSTD2 Enhancer +4.7kb gRNA 2 | GAAGACACTGAAGCTCATAG | AGG |
| TACSTD2 Enhancer +4.7kb gRNA 3 | AAGCCCTGGTGACCTACACA | AGG |
| TACSTD2 Enhancer +4.7kb gRNA 4 | CAGTGAACCAGTGAACTGTA | AGG |
| TACSTD2 Enhancer -7.1kb gRNA 1 | TGTAAGGGCTTTACACTCTG | GGG |
| TACSTD2 Enhancer -7.1kb gRNA 2 | AGCAGAACTGTACACATCAC | AGG |
| TACSTD2 Enhancer -7.1kb gRNA 3 | GCACCGTGACTCATCAGTGT | GGG |
| TACSTD2 Enhancer -7.1kb gRNA 4 | GATGTGTGATGCCCTGTTAA | GGG |
| TACSTD2 Enhancer -15.2kb gRNA 1 | TGGATTCAAGATCGAAACCC | TGG |
| TACSTD2 Enhancer -15.2kb gRNA 2 | TGACTTCATCAGATGGAAAG | TGG |
| TACSTD2 Enhancer -15.2kb gRNA 3 | GACATGAACGAGTCCTTGGA | AGG |
| TACSTD2 Enhancer -15.2kb gRNA 4 | TGGGCAGGCTGATTAATGAG | GGG |
| TGFA Enhancer +44kb gRNA 1 | TGAGTACCCAGCTCATGCA | AGG |

| | | |
|---|---|---|
| TGFA Enhancer +44kb gRNA 2 | GGGCATTACTCAGGCGCCC | GGG |
| TGFA Enhancer +44kb gRNA 3 | CCCTGGTTAAGAAGTTGTT | GGG |
| TGFA Enhancer +44kb gRNA 4 | GACAGGTGTTCTAAGCTAT | AGG |
| TGFA Promoter gRNA 1 | CCCCGGTCGCCGAGTGGCG | AGG |
| TGFA Promoter gRNA 2 | GAGCGCACCGAGCAGGGCG | CGG |
| TGFA Promoter gRNA 3 | GCACGGCGCTGGCAGTCGG | GGG |
| TGFA Promoter gRNA 4 | TAGGTAACGGGTGCCCGGG | CGG |
| TGFA Enhancer -43kb gRNA 1 | GGAAGCGGGATGCAGACCT | GGG |
| TGFA Enhancer -43kb gRNA 2 | CAAGACATCAGGCTTCCTG | GGG |
| TGFA Enhancer -43kb gRNA 3 | GTTCACCACAGCGTGTCAA | GGG |
| TGFA Enhancer -43kb gRNA 4 | CCAGGATCTGCCACCCCTG | TGG |
| TGFA Enhancer -53kb gRNA 1 | CACCGAATCAACTGAGGTC | AGG |
| TGFA Enhancer -53kb gRNA 2 | GGAGTTCCTACAATCACAG | AGG |
| TGFA Enhancer -53kb gRNA 3 | AAGACAACGGATGTGTCCG | TGG |
| TGFA Enhancer -53kb gRNA 4 | AGAAGCCCGAACCCAGAGA | AGG |
| TGFA Enhancer -54kb gRNA 1 | GAACGGGGACTTGAGGTAT | TGG |
| TGFA Enhancer -54kb gRNA 2 | AGCATCACGCTGAGAATGG | GGG |
| TGFA Enhancer -54kb gRNA 3 | GGACATCCACTTGAAAAGC | TGG |
| TGFA Enhancer -54kb gRNA 4 | TAAAATGAGTGCTCTAGGC | AGG |
| TGFA Enhancer -62kb gRNA 1 | AGGGCTACCTAACTGCAAT | AGG |
| TGFA Enhancer -62kb gRNA 2 | ATTGTCTGAACTCCGGATG | TGG |
| TGFA Enhancer -62kb gRNA 3 | TCAGGGAGTGTTGCAACCT | TGG |
| TGFA Enhancer -62kb gRNA 4 | GTATCACAGCGAAGCGGGT | GGG |

**Table S2. qPCR primers**

| Primer name | Sequence |
|---|---|
| TACSTD2_F | ACAACGATGGCCTCTACGAC |
| TACSTD2_R | GTCCAGGTCTGAGTGGTTGAA |
| PEG10_F | GAGCACCAGGGATTTCTCAGT |
| PEG10_R | GGTAGTTGTGCATCAGGTAGTG |
| TGFA_F | AGGTCCGAAAACACTGTGAGT |
| TGFA_R | AGCAAGCGGTTCTTCCCTTC |
| IL1RN_F | GGAATCCATGGAGGGAAGAT |
| IL1RN_R | TGTTCTCGCTCAGGTCAGTG |
| dCas9_F | GTGACCGAGGGAATGAGAAA |
| dCas9_R | AGCTGCTTCACGGTCACTTT |

**Table S3. Genomic Source Information**
* - indicates experiments conducted for this study. See Data Availability.

| Cell Line | Genomic Feature | Reference | Accession |
|---|---|---|---|
| Ishikawa | CTCF | (ENCODE, 2012) | GSE32465 |
| Ishikawa | Dnase | (ENCODE, 2012) | GSE32970 |
| Ishikawa | ER | (ENCODE, 2012; Rodriguez et al., 2020) | GSE32465; GSE129803 |
| Ishikawa | FOXA1 | (ENCODE, 2012) | GSE32465 |
| Ishikawa | GC Content | (Kent et al., 2002) | |
| Ishikawa | Gene length | (Kent *et al.*, 2002) | |
| Ishikawa | H3K27ac | (Vahrenkamp et al., 2018) | GSE109893 |
| Ishikawa | H3K27me3 | * | |
| Ishikawa | H3K36me3 | * | |
| Ishikawa | H3K4me1 | * | |
| Ishikawa | H3K4me3 | * | |
| Ishikawa | H3K9me3 | (Carleton et al., 2017) | GSE99906 |
| Ishikawa | JUN | * | |
| Ishikawa | LSD1 | * | |
| Ishikawa | MAX | * | |
| Ishikawa | MYC | * | |
| Ishikawa | Number Enhancers | * | |
| Ishikawa | Number ERBS | * | |
| Ishikawa | p300 | (ENCODE, 2012) | GSE32465 |
| Ishikawa | PRO-seq | * | |
| Ishikawa | RARA | * | |
| Ishikawa | RNAPII | (Carleton *et al.*, 2017; ENCODE, 2012) | GSE32465; GSE99906 |

| | | | |
|---|---|---|---|
| Ishikawa | SIN3A | * | |
| Ishikawa | TAF1 | * | |
| Ishikawa | scRNA-seq | * | |
| Ishikawa | HiChIP | * | |
| T-47D | CTCF | (ENCODE, 2012) | GSE32465 |
| T-47D | Dnase | (ENCODE, 2012) | GSE32970 |
| T-47D | ER | (ENCODE, 2012) | GSE32465 |
| T-47D | FOXA1 | (ENCODE, 2012) | GSE32465 |
| T-47D | GC Content | (Kent *et al.*, 2002) | |
| T-47D | Gene length | (Kent *et al.*, 2002) | |
| T-47D | H3K27ac | (Carleton et al., 2020; Zhang et al., 2016b) | GSE63109; GSE147141 |
| T-47D | H3K27me3 | * | |
| T-47D | H3K36me3 | (Zhang et al., 2016a) | GSE63109 |
| T-47D | H3K4me1 | * | |
| T-47D | H3K4me3 | * | |
| T-47D | H3K9me3 | (Zhang *et al.*, 2016a) | GSE63109 |
| T-47D | JUN | * | |
| T-47D | LSD1 | * | |
| T-47D | MAX | * | |
| T-47D | MYC | * | |
| T-47D | Number Enhancers | * | |
| T-47D | Number ERBS | * | |
| T-47D | p300 | (ENCODE, 2012) | GSE32465 |
| T-47D | PRO-seq | * | |
| T-47D | RARA | * | |
| T-47D | RNAPII | * | |
| T-47D | SIN3A | * | |
| T-47D | TAF1 | * | |
| T-47D | scRNA-seq | * | |
| T-47D | HiChIP | * | |
| LNCaP | H3ac | (Takayama et al., 2015; Yamamoto et al., 2019) | GSE62492 |
| LNCaP | AR | (Takayama *et al.*, 2015; Yamamoto *et al.*, 2019) | GSE62492 |
| LNCaP | CHD1 | (Metzger et al., 2016) | GSE64530 |
| LNCaP | ETV1 | (Chen et al., 2013) | GSE47120 |
| LNCaP | FOXP1 | (Takayama *et al.*, 2015; Yamamoto *et al.*, 2019) | GSE62492 |
| LNCaP | H3K27ac | (Taberlay et al., 2016) | GSE73785 |

| | | | |
|---|---|---|---|
| LNCaP | H3K27me3 | (Takayama *et al.*, 2015; Yamamoto *et al.*, 2019) | GSE62492 |
| LNCaP | H3K4me1 | (Takayama *et al.*, 2015; Yamamoto *et al.*, 2019) | GSE62492 |
| LNCaP | H3K4me3 | (Takayama *et al.*, 2015; Yamamoto *et al.*, 2019) | GSE62492 |
| LNCaP | LSD1 | (Metzger *et al.*, 2016) | GSE64530 |
| LNCaP | RUNX1 | (Takayama *et al.*, 2015; Yamamoto *et al.*, 2019) | GSE62492 |
| LNCaP | scRNA-seq | (Taavitsainen et al., 2021) | GSE168668 |
| LNCaP | HiChIP | * | |

**Supplemental References**


Carleton, J.B., Berrett, K.C., and Gertz, J. (2017). Multiplex Enhancer Interference Reveals Collaborative Control of Gene Regulation by Estrogen Receptor α-Bound Enhancers. Cell Syst *5*, 333-344.e335. 10.1016/j.cels.2017.08.011.

Carleton, J.B., Ginley-Hidinger, M., Berrett, K.C., Layer, R.M., Quinlan, A.R., and Gertz, J. (2020). Regulatory sharing between estrogen receptor α bound enhancers. Nucleic Acids Research *48*, 6597-6610. 10.1093/nar/gkaa454.

Chen, Y., Chi, P., Rockowitz, S., Iaquinta, P.J., Shamu, T., Shukla, S., Gao, D., Sirota, I., Carver, B.S., and Wongvipat, J. (2013). ETS factors reprogram the androgen receptor cistrome and prime prostate tumorigenesis in response to PTEN loss. Nature medicine *19*, 1023-1029.

ENCODE, C. (2012). An integrated encyclopedia of DNA elements in the human genome. Nature *489*, 57-74. 10.1038/nature11247.

Ginley-Hidinger, M., Carleton, J.B., Rodriguez, A.C., Berrett, K.C., and Gertz, J. (2019). Sufficiency analysis of estrogen responsive enhancers using synthetic activators. Life Science Alliance *2*, e201900497. 10.26508/lsa.201900497.

Kent, W.J., Sugnet, C.W., Furey, T.S., Roskin, K.M., Pringle, T.H., Zahler, A.M., and Haussler, D. (2002). The human genome browser at UCSC. Genome research *12*, 996-1006.

Metzger, E., Willmann, D., McMillan, J., Forne, I., Metzger, P., Gerhardt, S., Petroll, K., Von Maessenhausen, A., Urban, S., and Schott, A.-K. (2016). Assembly of methylated KDM1A and CHD1 drives androgen receptor–dependent transcription and translocation. Nature structural & molecular biology *23*, 132-139.

Rodriguez, A.C., Vahrenkamp, J.M., Berrett, K.C., Clark, K.A., Guillen, K.P., Scherer, S.D., Yang, C.-H., Welm, B.E., Janát-Amsbury, M.M., and Graves, B.J. (2020). ETV4 is necessary for estrogen signaling and growth in endometrial cancer cells. Cancer research *80*, 1234-1245.

Taavitsainen, S., Engedal, N., Cao, S., Handle, F., Erickson, A., Prekovic, S., Wetterskog, D., Tolonen, T., Vuorinen, E.M., Kiviaho, A., et al. (2021). Single-cell ATAC and RNA sequencing reveal pre-existing and persistent cells associated with prostate cancer relapse. Nat Commun *12*, 5307. 10.1038/s41467-021-25624-1.

Taberlay, P.C., Achinger-Kawecka, J., Lun, A.T., Buske, F.A., Sabir, K., Gould, C.M., Zotenko, E., Bert, S.A., Giles, K.A., and Bauer, D.C. (2016). Three-dimensional disorganization of the cancer genome occurs coincident with long-range genetic and epigenetic alterations. Genome research *26*, 719-731.

Takayama, K., Suzuki, T., Tsutsumi, S., Fujimura, T., Urano, T., Takahashi, S., Homma, Y., Aburatani, H., and Inoue, S. (2015). RUNX1, an androgen- and EZH2-regulated gene, has differential roles in AR-dependent and -independent prostate cancer. Oncotarget *6*, 2263-2276. 10.18632/oncotarget.2949.

Vahrenkamp, J.M., Yang, C.H., Rodriguez, A.C., Almomen, A., Berrett, K.C., Trujillo, A.N., Guillen, K.P., Welm, B.E., Jarboe, E.A., Janat-Amsbury, M.M., and Gertz, J. (2018). Clinical and Genomic Crosstalk between Glucocorticoid Receptor and Estrogen Receptor α In Endometrial Cancer. Cell Rep *22*, 2995-3005. 10.1016/j.celrep.2018.02.076.

Yamamoto, S., Takayama, K.i., Obinata, D., Fujiwara, K., Ashikari, D., Takahashi, S., and Inoue, S. (2019). Identification of new octamer transcription factor 1-target genes upregulated in castration-resistant prostate cancer. Cancer Science *110*, 3476-3485.

Zhang, G., Zhao, Y., Liu, Y., Kao, L.-P., Wang, X., Skerry, B., and Li, Z. (2016a). FOXA1 defines cancer cell specificity. Science Advances *2*, e1501473. doi:10.1126/sciadv.1501473.

Zhang, G., Zhao, Y., Liu, Y., Kao, L.P., Wang, X., Skerry, B., and Li, Z. (2016b). FOXA1 defines cancer cell specificity. Sci Adv *2*, e1501473. 10.1126/sciadv.1501473.
